# Associations between common genetic variants and income provide insights about the socioeconomic health gradient

**DOI:** 10.1101/2024.01.09.574865

**Authors:** Hyeokmoon Kweon, Casper A.P. Burik, Yuchen Ning, Rafael Ahlskog, Charley Xia, Erik Abner, Yanchun Bao, Laxmi Bhatta, Tariq O. Faquih, Maud de Feijter, Paul Fisher, Andrea Gelemanović, Alexandros Giannelis, Jouke-Jan Hottenga, Bita Khalili, Yunsung Lee, Ruifang Li-Gao, Jaan Masso, Ronny Myhre, Teemu Palviainen, Cornelius A. Rietveld, Alexander Teumer, Renske M. Verweij, Emily A. Willoughby, Esben Agerbo, Sven Bergmann, Dorret I. Boomsma, Anders D. Børglum, Ben M. Brumpton, Neil Martin Davies, Tõnu Esko, Scott D. Gordon, Georg Homuth, M. Arfan Ikram, Magnus Johannesson, Jaakko Kaprio, Michael P. Kidd, Zoltán Kutalik, Alex S.F. Kwong, James J. Lee, Annemarie I. Luik, Per Magnus, Pedro Marques-Vidal, Nicholas G. Martin, Dennis O. Mook-Kanamori, Preben Bo Mortensen, Sven Oskarsson, Emil M. Pedersen, Ozren Polašek, Frits R. Rosendaal, Melissa C. Smart, Harold Snieder, Peter J. van der Most, Peter Vollenweider, Henry Völzke, Gonneke Willemsen, Jonathan P. Beauchamp, Thomas A. DiPrete, Richard Karlsson Linnér, Qiongshi Lu, Tim T. Morris, Aysu Okbay, K. Paige Harden, Abdel Abdellaoui, W. David Hill, Ronald de Vlaming, Daniel J. Benjamin, Philipp D. Koellinger

## Abstract

We conducted a genome-wide association study (GWAS) on income among individuals of European descent and leveraged the results to investigate the socio-economic health gradient (*N*=668,288). We found 162 genomic loci associated with a common genetic factor underlying various income measures, all with small effect sizes. Our GWAS-derived polygenic index captures 1 - 4% of income variance, with only one-fourth attributed to direct genetic effects. A phenome-wide association study using this polygenic index showed reduced risks for a broad spectrum of diseases, including hypertension, obesity, type 2 diabetes, coronary atherosclerosis, depression, asthma, and back pain. The income factor showed a substantial genetic correlation (0.92, *s.e*. = .006) with educational attainment (EA). Accounting for EA’s genetic overlap with income revealed that the remaining genetic signal for higher income related to better mental health but reduced physical health benefits and increased participation in risky behaviours such as drinking and smoking.

## Introduction

Income is a crucial determinant of individuals’ access to resources and overall quality of life. Extensive evidence shows that higher income is positively correlated with increased subjective well-being, better health, and longer life expectancy.^1–5^ For instance, the gap in life expectancy between the richest and poorest 1% of individuals in the US has been estimated to be 14.6 years for men (95% CI, 14.4 to 14.8 years) and 10.1 years for women (95% CI, 9.9 to 10.3 years).^6^ Notably, higher income is associated with increased longevity and well-being across the entire income distribution, highlighting its broad relevance in current society.^3,6,7^

Income is a complex phenotype influenced by many factors, including environmental conditions and education.^8,9^ Parents’ socio-economic status shapes a child’s developmental trajectory, including their skills, behaviours, educational attainment, career prospects, and eventual adult income.^10,11^ Moreover, certain heritable individual characteristics, such as cognitive ability and personality traits,^12–14^ are well-known predictors of income within contemporary Western societies. Twin studies have estimated income heritability in these societies to be around 40-50%.^15–17^ However, the heritability of income and its associated genes are not fixed; rather, they reflect social realities shaped by technological, institutional, and cultural factors.^18^ These factors are malleable and exhibit variations across different regions and historical epochs, which can lead to fluctuations in heritability estimates for socio-economic status (SES) over time^19,20^ and imperfect genetic correlations across samples.^21^

The results from statistically well-powered GWAS of SES present numerous opportunities to shed light on these social realities. For example, they allow investigating questions about sex differences in labour market processes, cross-country comparisons in the genetic architecture of income, and investigating the processes contributing to intergenerational social mobility.^22^ They also facilitate studies investigating the interaction effects between genetic and environmental factors. Furthermore, they enable the exploration of genetic correlations between income and health outcomes, potentially unveiling new insights into the socioeconomic health gradient.

Two previous GWAS have been conducted on household income.^23,24^ The first was in a sample of 96,900 participants from the initial release of the UK Biobank (UKB)^25^ and found two loci. The second was carried out in the full release of the UKB with 286,301 individuals and found 30 approximately uncorrelated loci. A meta-analysis of these results with the genetically correlated trait educational attainment increased the effective sample size to 505,541 individuals and identified 144 loci. A recent GWAS on occupational status in the UKB data identified cognitive skills, scholastic motivation, occupational aspiration, personality traits, and behavioural disinhibition (proxied by ADHD) as potential mediating factors linking genetics to occupational status.^26^

Building on these earlier contributions, we conducted a GWAS leveraging multiple income measures. We ran sex-stratified analyses and meta-analyzed results from 32 cohorts across 12 economically advanced countries and three continents, yielding the largest GWAS on income to date with an effective sample size of *N* = 668,288 (Table 1). Due to data availability and statistical power considerations, our analyses and conclusions are restricted to individuals carrying genotypes most similar to the EUR panel of the 1000 Genomes dataset, as compared to individuals sampled elsewhere in the world (1KG-EUR-like individuals).

**Table 1.**
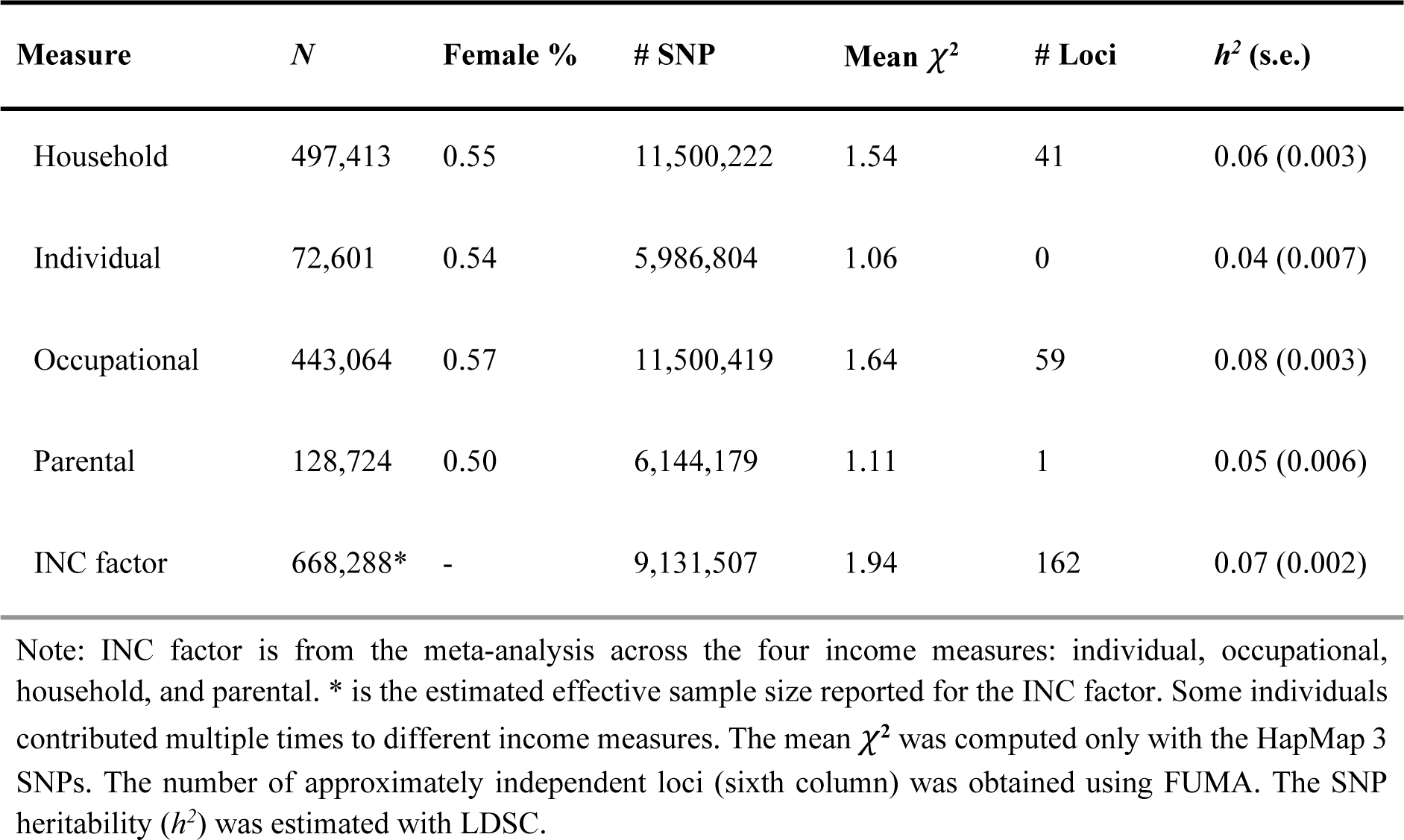
GWAS summary.

The greater statistical power of our GWAS enabled us to conduct a series of follow-up analyses that investigate the socio-economic health gradient from a genetic perspective. In particular, we leveraged the data to compare the GWAS results for income and EA to disentangle their unique genetic correlates with health. Furthermore, our multi-sample approach and sex-specific GWAS results allowed us to test for possible differences in the genetic architecture of income across samples and sexes.

For a less technical description of the paper and how it should -- and should not -- be interpreted, see the **Frequently Asked Questions** document (FAQ) and **Box 1**.

## Results

### Multivariate GWAS of income

We used four measures of income (individual, occupational, household, and parental income) and conducted a GWAS meta-analysis of their shared genetic basis. **Supplementary Information section 2.1** discusses the differences between these measures and their relative advantages and disadvantages as proxies for individual income. Dropping parental income from the meta-analysis leads to a slight statistical power decrease but does not qualitatively change our results.

A sex-stratified GWAS was carried out on each available income measure in each cohort. We restricted our analyses to 1KG-EUR-like individuals who were not currently enrolled in an educational program or who were aged above 30 if their current enrollment status was unknown. The natural log transformation was applied to the income measures. We applied standardised quality control procedures to each cohort-level result (see **Supplementary Information section 2.4** for details). For each sex and each income measure, we performed a sample-size-weighted meta-analysis with METAL.^27^ We then meta-analyzed the male and female results of each income measure using MTAG,^28^ which accounts for any potential genetic relatedness between the male and female samples.

Across the four GWAS on different income measures, we identified 86 non-overlapping loci in the genome (see Supplementary Information section 2.6 for the definition of loci and lead SNPs). **Table 1** summarises the results. Occupational and household income produced the most genetic associations (59 and 41 loci, respectively), as expected based on sample sizes and SNP-based heritability estimates based on linkage disequilibrium score regression (LDSC) (*h*^2^ = 0.08 (*s.e.* = 0.003) and 0.06 (*s.e.* = 0.003), respectively). The four income measures’ pairwise genetic correlation (*r_g_*) estimates demonstrated substantial shared genetic variance, with all pairwise *r_g_*’s at least 0.8 (**Fig. 1a**).

**Fig. 1.**
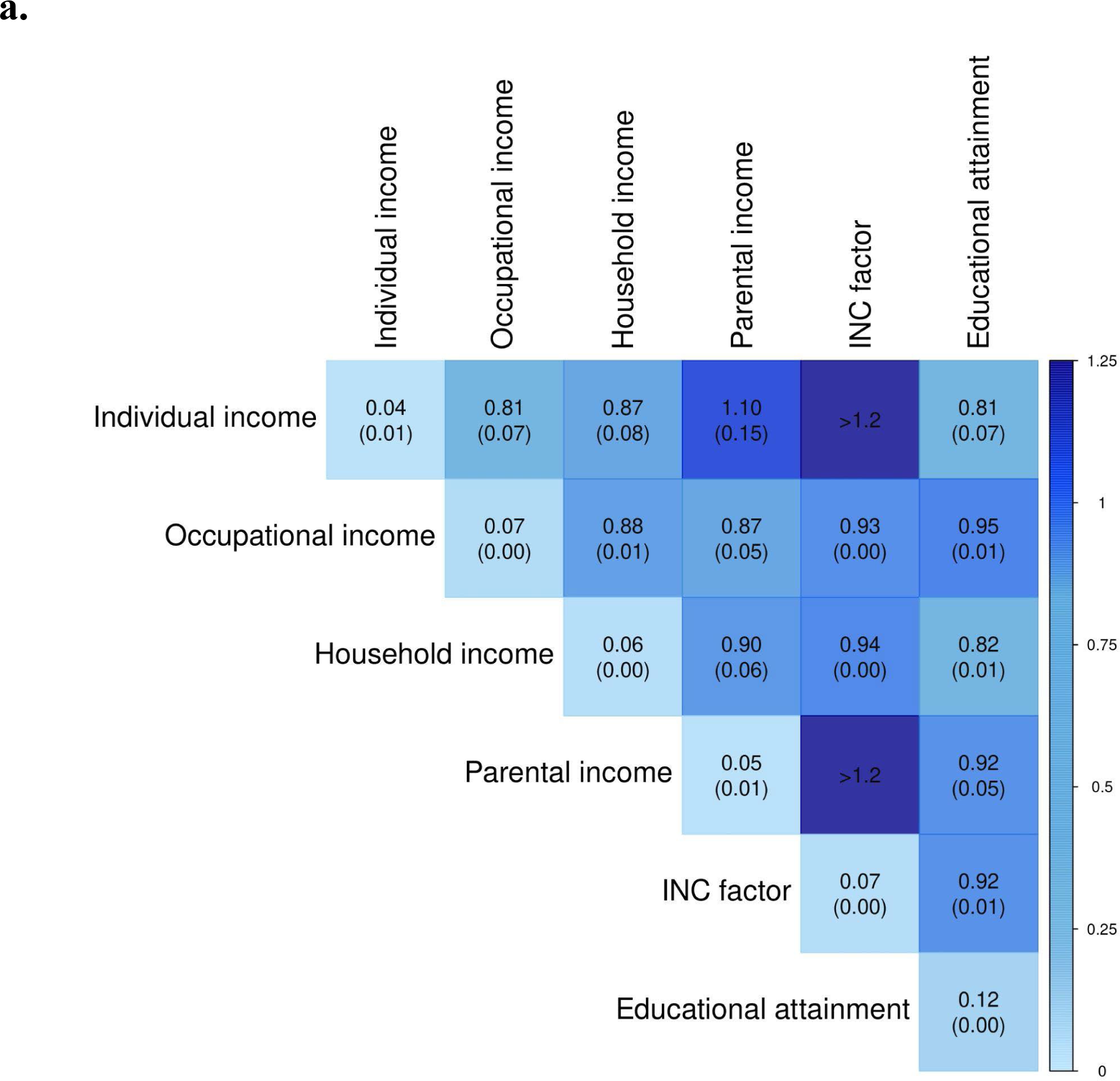
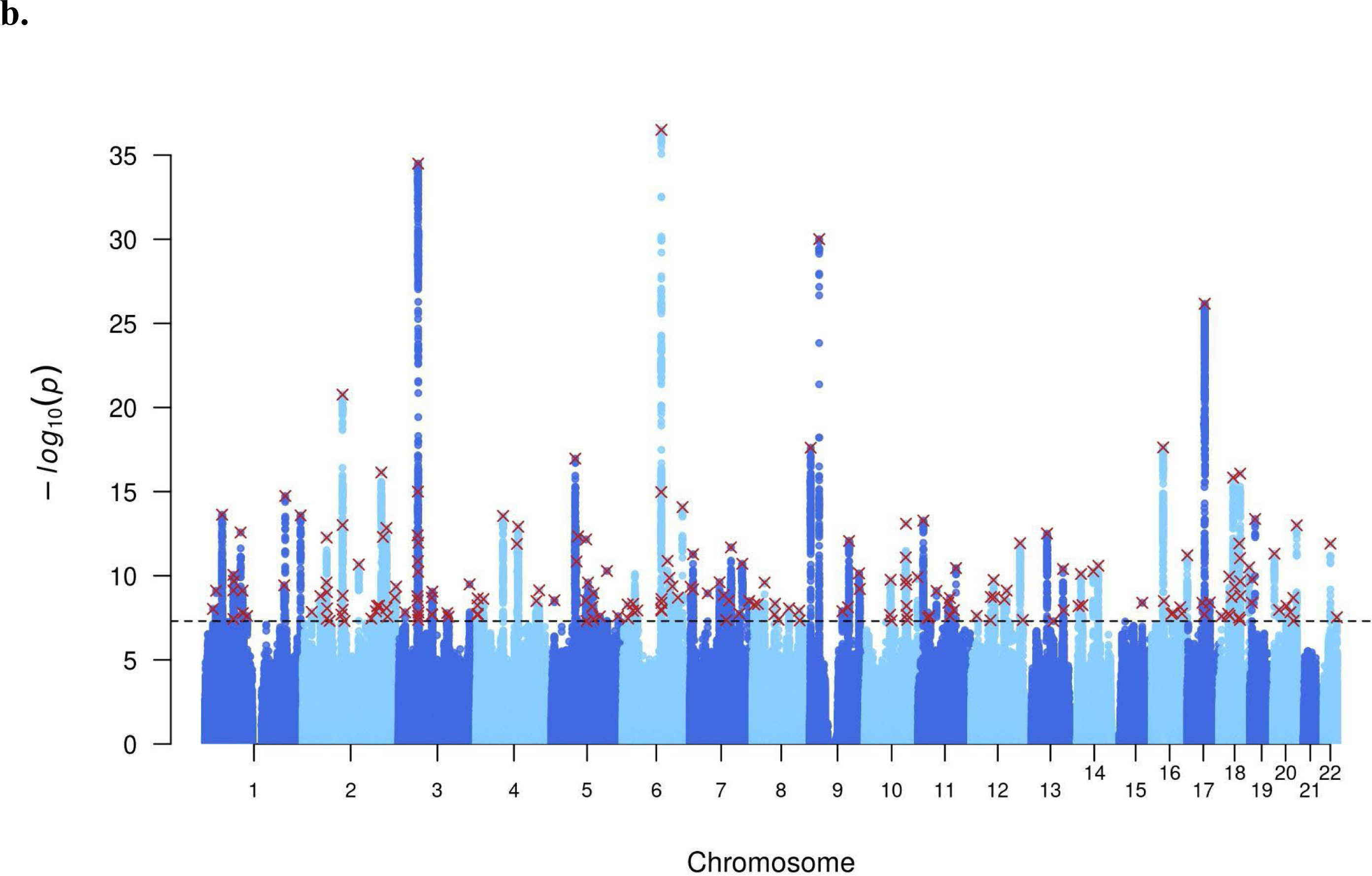
Multivariate genome-wide association study of income. **a.** LD score regression (LDSC) estimates of pairwise genetic correlations between the four input income measures, the meta-analyzed income (INC factor), and educational attainment. The diagonal elements report SNP heritabilities from LDSC. The standard errors are reported in the parentheses. Some of the results were out-of-bound estimates (exceeding 1.2). **b.** Manhattan plot presenting the GWAS results of INC factor. *P* values are plotted on -*log_10_* scale. The red crosses indicate the lead SNPs found from FUMA (*r*^2^ < 0.1).

Next, we meta-analyzed the association results across the four income measures using MTAG (see Supplementary Information section 2.5 for details). We observed that the MTAG procedure yields nearly identical results to genomic SEM’s common factor function.^29^ Thus, we hereafter refer to the meta-analyzed income as ‘the income factor’ (INC factor). Since MTAG already applies a bias correction with the intercept from LDSC,^30^ we did not apply further adjustments for cryptic relatedness and population stratification.

The INC factor GWAS was estimated to have an effective sample size of 668,288, based on occupational income’s heritability scale (*N_eff_* = 1,198,347 based on individual income). The meta-analysis across the income measures led to a substantial increase in power, which allowed us to identify 162 loci tagged by 207 lead SNPs (**Fig. 1b**). 88 of these loci were newly identified compared to the previously published GWAS household income result conducted in the UKB.^24^ The genetic correlation of the previous household income GWAS result was 0.92 (*s.e.* = 0.008) with the INC factor and 0.94 (*s.e.* = 0.006) when we restrict our analysis to only our household income measure.

The effect sizes of the lead SNPs were small. Adjusting for the statistical winner’s curse, one additional count in the effect allele of the median lead SNP was associated with an increase in income of 0.30% (these effect-size calculations require an assumption about the standard deviation of the dependent variable because MTAG yields standardised effect-size estimates; we use the standard deviation estimate of log hourly occupational wage from the UKB, which is 0.35). The estimated effects at the 5^th^ and 95^th^ percentiles were 0.18 and 0.60%, respectively (see **Supplementary Information section 2.7**). To put these estimates into perspective, the median annual earnings of full-time workers in the US was $56,473 in 2021.^31^ A 0.3% increase would equal an additional annual income of $169. In terms of the variance explained (*R*^2^), all of the lead SNPs each had *R*^2^ lower than 0.011% after adjustment for the statistical winner’s curse (**Supplementary Fig. 2**).

### Cross-sex and cross-country heterogeneity

The heritability of income and its genetic associations may vary across different social environments or different groups within an environment. To investigate the potential heterogeneity of genetic associations with income, we examined cross-cohort genetic correlations. We found that the inverse-variance weighted mean genetic correlations across pairs of cohorts were 0.45 (*s.e.* = 0.22) for individual income, 0.52 (*s.e.* = 0.13) for household income, and 0.90 (*s.e.* = 0.24) for occupational income (**Supplementary Tables 28a-c**).

Next, we meta-analyzed cohorts from the same country with the same income measure available and estimated the genetic correlations across these countries (Estonia, Netherlands, Norway, United Kingdom, USA - **Extended Data Figure 1a**). For most country-pairs, the genetic correlation of the same income measure is >0.8. While meta-analysis increases statistical power and yields more precise estimates of the average effect size, it also tends to mask non-random heterogeneity in effect size estimates across samples. Despite this latter point, we find that occupational income in Norway displayed lower genetic correlations with occupational or household income in other countries, ranging from 0.43 (*s.e.* = 0.23) to 0.82 (*s.e.* = 0.10). Similarly, occupational income’s genetic correlation with educational attainment (EA) was also lower in Norway (*r_g_* = 0.69, *s.e.* = 0.08) compared to the other countries. These findings align with phenotypic evidence that ranks Norway lowest among OECD countries in terms of financial returns for obtaining a college degree.^32^ Next, we investigated whether the large number of samples from the United Kingdom in our meta-analysis could have skewed our results. To address this, we conducted a separate meta-analysis procedure for the British and non-British cohorts, comprising participants from 10 countries. We obtained two distinct GWAS results for the INC factor and found a perfect genetic correlation of 1.001 (*s.e.* = 0.03) between them. Thus, the average effect sizes of SNPs associated with the INC factor are almost identical in British and non-British cohorts.

We observed slight between-sex heterogeneity in the genetic associations of income, as supported by the evidence presented in **Extended Data Figure 1b**. The estimated between-sex genetic correlations based on meta-analysed GWAS results for individual, occupational, and household income were 1.06 (s.e. = 0.32), 0.91 (s.e. = 0.03), and 0.95 (s.e. = 0.03), respectively. Notably, the latter two estimates were statistically distinguishable from unity but remained above 0.9. Most cohort-specific cross-sex genetic correlations for income are too noisy to be interpreted (**Supplementary Tables 17b-d**). One exception is the UK Biobank sample, which shows a non-perfect genetic correlation between men and women for occupational income (*r*_g_ = 0.91, *s.e.* = 0.03). Another exception is the Danish iPsych cohort, where we estimated a genetic correlation of 0.76 (s.e. = 0.10) between maternal and paternal income. These findings are consistent with the hypothesis that men and women face non-identical labour market conditions. The lower genetic correlation between maternal and paternal income suggests that differences in labour market conditions were more pronounced in previous generations.

We also conducted the INC factor GWAS for the male and female results separately and found that their genetic correlation was statistically indistinguishable from one (*r_g_* = 0.98, *s.e.* = 0.02).

### Comparison with educational attainment

To compare the GWAS results for the INC factor with those for EA, we first conducted an auxiliary GWAS on EA to obtain the most-powered GWAS result of EA with the summary statistics currently available to us: We first carried out a GWAS of EA in the UKB, based on the protocol of the latest EA GWAS (EA4).^33^ We then meta-analyzed these GWAS results with the EA3 summary statistics^21^ that did not include the UKB, using the meta-analysis version of MTAG. While previous GWASs on income found somewhat inconsistent results on the genetic correlation between educational attainment (EA)^21,33^ and income (*r_g_* = 0.90 (*s.e.* = 0.04)^23^ and 0.77 (*s.e.* = 0.02)^24^), with much greater precision, we found a high genetic correlation that is very close to the first reported estimate (*r_g_* = 0.917, *s.e.* = 0.006). Among the input income measures, the genetic correlation with EA was higher for occupational and parental income (*r_g_* = 0.95 and 0.92; *s.e. =* 0.01 and 0.05 respectively) and lower for individual and household income (*r_g_* = 0.81 and 0.82; *s.e. =* 0.07 and 0.01 respectively). Furthermore, 138 out of 161 loci for the INC factor overlapped with those for EA.

The *r_g_* estimate of 0.917 between the INC factor and EA implies that only 1 - 0.917^2^ = ∼16% of the genetic variance of the INC factor would remain once the genetic covariance with EA was statistically removed. We employed the GWAS-by-subtraction approach using Genomic SEM^29^ to identify this residual genetic signal (referred to as ‘NonEA-INC’). We identified one locus of genome-wide significance for NonEA-INC, marked by the lead SNP rs34177108 on chromosome 16 (**Extended Data Fig 2c**). This locus was previously found to be associated with vitamin D levels and hair and skin-related traits such as colour, sun exposure, and cancer, possibly picking up on uncontrolled population stratification or physical traits linked to discrimination in the labour market.

### Polygenic prediction

We conducted polygenic index (PGI) analyses with individuals of European ancestry in the Swedish Twin Registry (STR), which was not included in our meta-analysis. We chose STR as the main prediction cohort because it has twins and administrative data on individual, occupational, and household income. In addition, we also used the UKB siblings (UKB-sib) and the Health and Retirement Study (HRS) from the US as prediction cohorts. For the UKB-sib, occupational and household income measures were available. For the HRS, a self-reported individual income measure was available. In the STR and the UKB-sib cohorts, except when examining within-family prediction, we randomly selected only one individual from each family.

After generating hold-out versions of GWAS on the INC factor and EA to remove the sample overlap with each prediction sample, we constructed PGIs for the INC factor and EA using LDpred2^34^. Before conducting prediction analyses, we residualised the log of income on demographic covariates, including a third-degree polynomial in age, year of observation, and interactions with sex. We measured the prediction accuracy as the incremental *R*^2^ from adding the PGI to a regression of the phenotype on a set of baseline covariates, which were the top 20 genetic principal components and genotype batch indicators.

A cohort-specific upper bound for the theoretically possible predictive accuracy of PGIs on income can be obtained by the GREML^35^ estimate of the SNP-based heritability of income, which is close to 10% for the available income measures in the STR and UKB sibling sample (**SI Table 13**).

The actual prediction accuracy of PGIs for income is lower than the theoretical maximum, primarily due to finite GWAS sample size but also due to imperfect genetic correlations across meta-analysed cohorts and differences in measurement accuracy of income across samples.^36^ In the STR (**Fig. 2**), the INC factor PGI predicted *ΔR*^2^ = 1.3% (95% CI: 1.0-1.6) for individual income, 3.7% (95% CI: 3.1-4.2) for occupational income, and 1.0% (95% CI: 0.6-1.4) for household income. The EA PGI had predictive accuracy results in a similar range for individual and household income, except for occupational income, for which the accuracy was larger: *ΔR*^2^ = 4.7% (95% CI: 4.0-5.4). In the UKB-sib, the predictive accuracy of the INC factor PGI was *ΔR*^2^ = 4.7% (95% CI: 4.3-5.2) for occupational income and 3.9% (95% CI: 3.5-4.3) for household income. The EA PGI achieved a better predictive accuracy for occupational income (*ΔR*^2^ = 6.9%, 95% CI: 6.3-7.4), while only slightly higher for household income (*ΔR*^2^ = 4.4%, 95% CI: 3.9-4.8). In the HRS, INC factor PGI had *ΔR*^2^ = 2.7% (95% CI: 2.1-3.3) for predicting individual income, which was close to the EA PGI’s result (*ΔR*^2^ = 3.1%, 95% CI: 2.4-3.8).

**Fig. 2.**
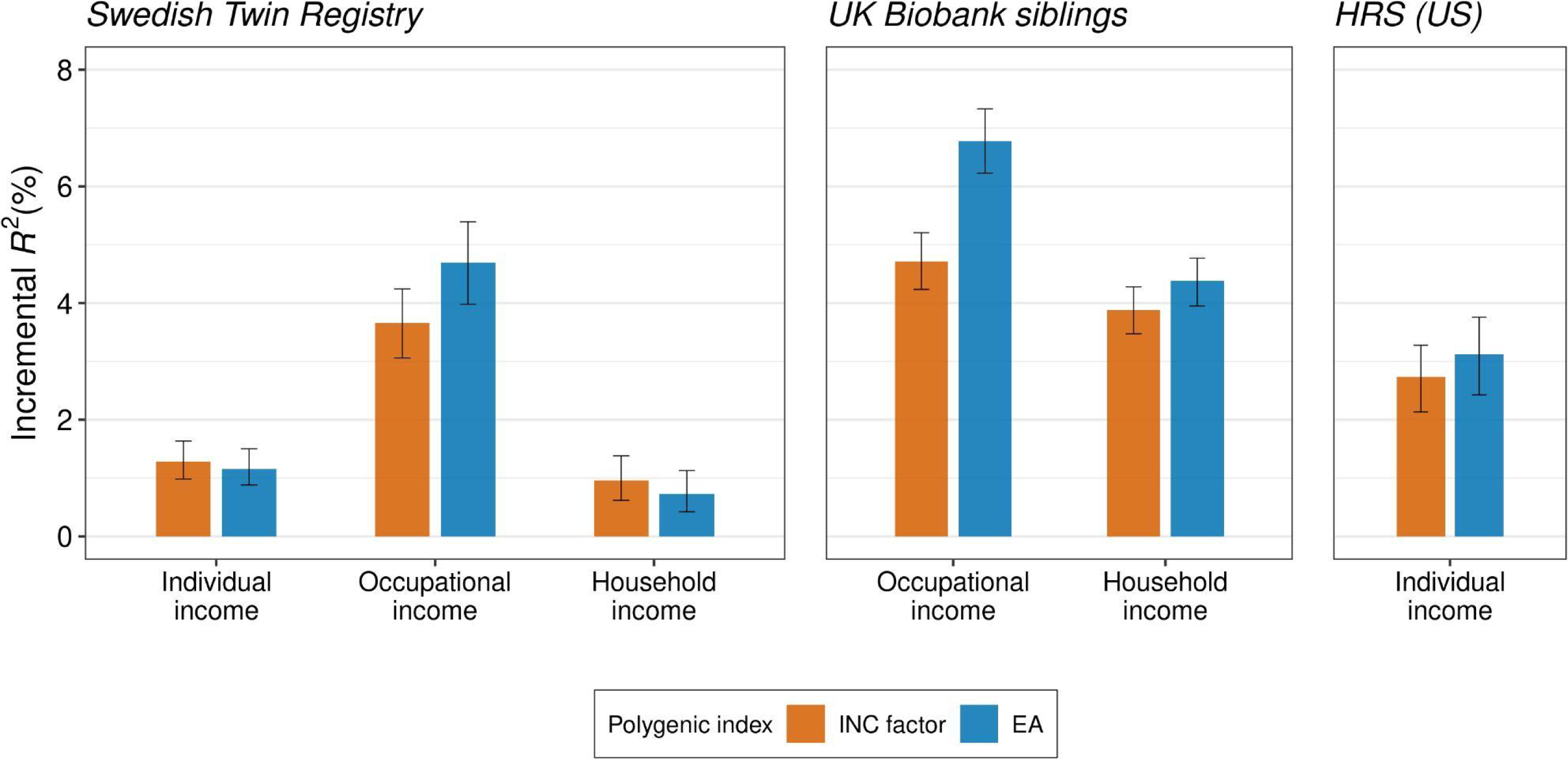
Polygenic prediction of income measures. The figure reports polygenic prediction results in the Swedish Twin Registry (STR), the UK Biobank (UKB) siblings, and the Health and Retirement Study (HRS) with polygenic indexes (PGI) for INC factor and EA. Prior to fitting the regressions, each phenotype was residualized of demographic covariates (sex, a third-degree polynomial in age, and interactions with sex) within each wave and the mean of the residuals was obtained across the waves for each individual (only a single wave for the UKB siblings). Incremental *R*^2^ is the difference between the *R*^2^ from regressing the residualized outcome on the PGI and the controls (20 genetic PCs and genotyping batch indicators) and the *R*^2^ from a regression only on the controls. Only individuals of European ancestry were included and one sibling from each family was randomly chosen: *N* = 24,946 (individual), 19,245 (occupational), and 15,655 (household) for the STR; 15,556 (occupational), and 18,303 (household) for the UKB siblings; and 6,171 (individual) for the HRS. The error bars indicate 95% confidence intervals obtained by bootstrapping the sample 1,000 times.

In terms of the coefficient estimates in the UKB-sib, one standard deviation increase in the INC factor PGI was associated with a 7.2% increase in the occupational income (95% CI: 6.7-7.7) and a 12.3% increase in the household income (95% CI: 11.4-13.2). These estimates were comparable to the effect of one additional year of schooling on income, whose estimates tend to range from 5 to 15%.^8,9,37^

The predictive power of the INC factor PGI approached zero once EA or the EA PGI was controlled for. In the UKB-sib, *ΔR*^2^ decreased below 1% for occupational and household income, while the estimates were still statistically different from zero (**Extended Data Fig. 3** and **Supplementary Table 21**).

We estimated the share of the direct genetic effect in the overall population effect captured by the INC factor PGI, following the recent approach that imputes parental genotypes from first-degree relatives.^33,38^ Using the UKB-sib sample, we isolated the direct effect of the PGI from the population effect on occupational and household income by controlling for parental PGIs. We found that the ratio of direct-to-population effect estimates is 0.51 (*s.e.* = 0.05) and 0.49 (*s.e.* = 0.05) for occupational and household income, respectively (**Supplementary Table 22**). These results imply that only 24.0% or 25.7% of the INC factor PGI’s predictive power was due to direct genetic effects, which was very close to the result for the EA PGI estimated elsewhere (25.5%).^38^

We next conducted a phenome-wide association study of the INC factor PGI based on electronic health records in the UKB siblings holdout sample. We tested 115 diseases with sex-specific sample prevalence no lower than 1%. In total, 50 diseases from different categories were associated with the INC factor PGI after Bonferroni correction and 14 after controlling for parental PGI (**Fig. 3, Extended Data Fig. 4** and **Supplementary Tables 27a-b**). In all cases, a higher INC factor PGI value was associated with reduced disease risk, including reduced risk for hypertension, gastroesophageal reflux disease (GERD), type 2 diabetes, obesity, osteoarthritis, back pain, and depression. The strongest association of a higher INC factor PGI and a disease was found for essential hypertension.

**Fig. 3.**
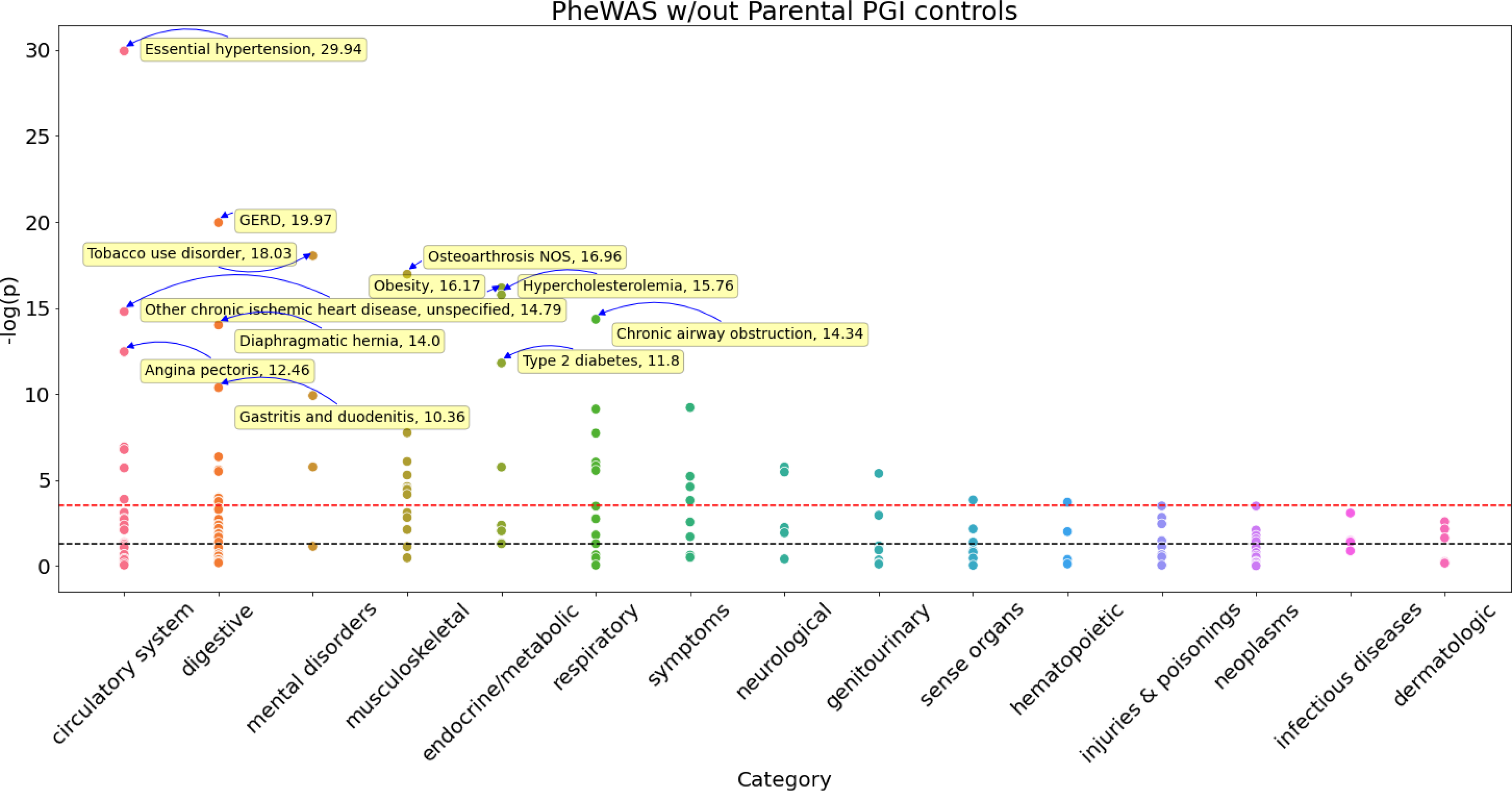
Phenome-wide association study of the INC factor PGI (without parental PGI controls) in electronic health records for the UKB sibling sample. This figure illustrates the genetic association of the INC factor PGI with 115 diseases from 15 categories without controlling for parental PGIs. The yellow boxes, with arrows pointing to the observations and -log10(p) values reported after the phenotypes, highlight diseases that are most strongly associated with the INC factor PGI (-log10(*p*)>10).

### Genetic correlations

We next explored the genetic correlations of the income (INC) factor, educational attainment (EA), and NonEA-INC with phenotypes related to behaviours, psychiatric disorders, and physical health (**Fig. 4**). LDSC estimates revealed that the genetic correlations of EA and the INC factor largely align. However, noticeable differences emerged for traits in the psychiatric and psychological domains. Specifically, NonEA-INC is associated with a reduced risk for certain psychiatric disorders previously reported to correlate positively with EA.^39–41^ These discrepancies were observed for schizophrenia (*r_g_* = -0.29, *s.e.* = 0.04), autism spectrum (*r_g_* = -0.27, *s.e.* = 0.06), and obsessive-compulsive disorder (*r_g_* = -0.22, *s.e.* = 0.08). One possible interpretation of these findings is that these psychiatric disorders have more severe negative effects on individual performance in the labour market than in the educational system.

**Fig. 4.**
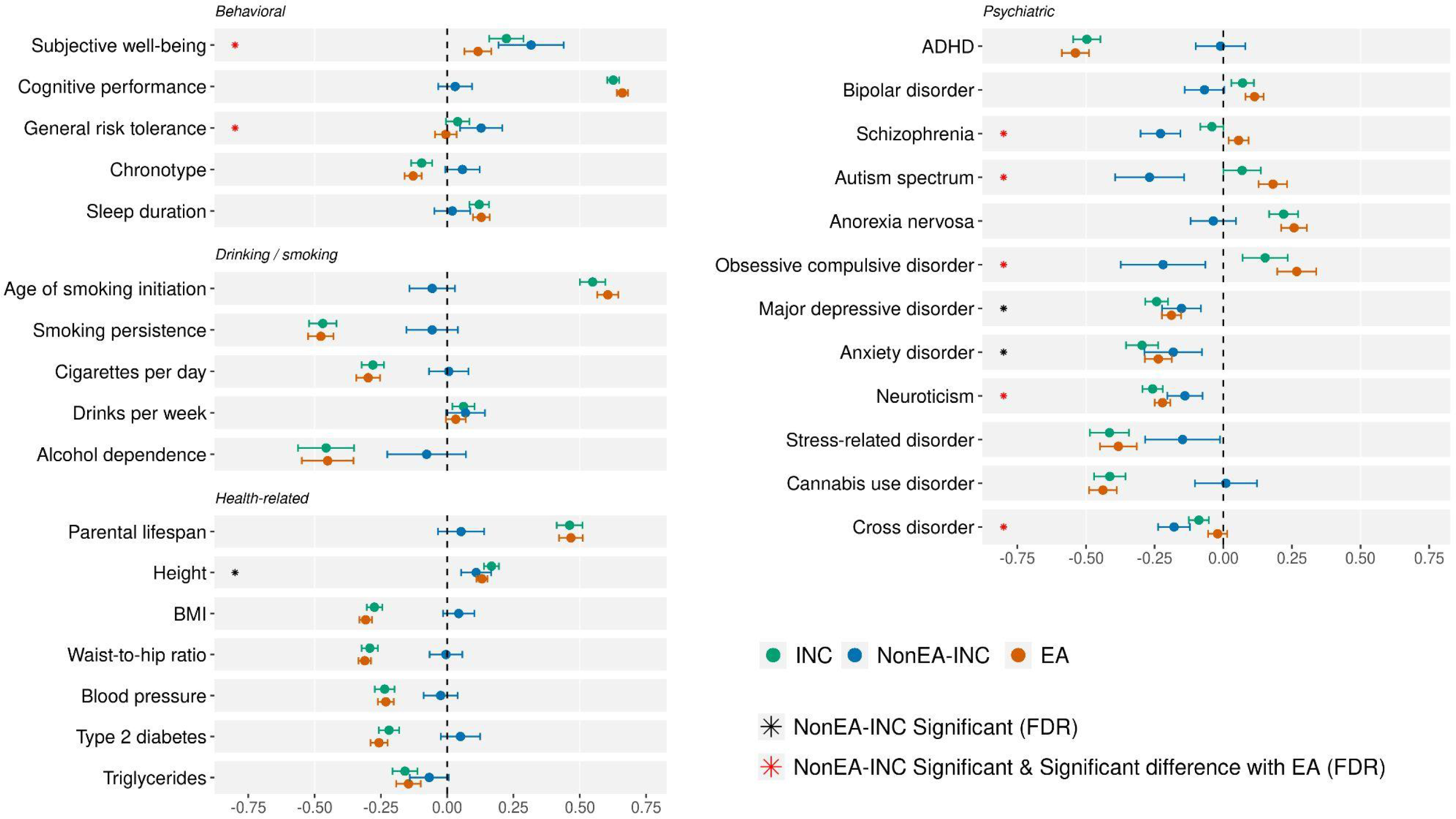
Genetic correlation estimates. Genetic correlation estimates of INC factor, NonEA-INC, and EA. The estimates were obtained from LDSC. The black asterisks indicate the statistical significance for NonEA-INC at the false discovery rate (FDR) of 5% and the asterisks were indicated in red if the estimate was also significantly different from the estimate for EA at the FDR of 5%. The standard error for the difference was computed from jackknife estimates.

Intriguingly, NonEA-INC exhibits a near-zero genetic correlation with cognitive performance (*r_g_* = 0.03, *s.e.* = 0.03). At the same time, both EA and the general income (INC) factor display strong positive genetic correlations with it (*r_g_* = 0.66, *s.e.* = 0.01 and *r_g_* = 0.63, *s.e.* = 0.01, respectively). This may suggest that high cognitive performance primarily influences income through education. Furthermore, this result is consistent with high income being attainable through social connections, inherited wealth, entrepreneurial activities, or the pursuit of well-paying jobs that do not require high cognitive performance.

While EA and the general INC factor have substantial negative genetic correlations with health-related behaviours such as age of smoking initiation, smoking persistence, cigarettes per day, and alcohol dependence, we found that NonEA-INC has near-zero genetic correlations with these traits (albeit the latter have substantially larger error margins of the point estimates).

NonEA-INC also displayed genetic correlations with other phenotypes that are similar to EA. Specifically, NonEA-INC had negative genetic correlations with major depressive disorder (*r_g_* = -0.15, *s.e.* = 0.04), anxiety disorder (*r_g_* = -0.19, *s.e.* = 0.05), and the related trait of neuroticism (*r_g_* = -0.14, *s.e.* = 0.03), but positive genetic correlations with subjective well-being (*r_g_* = 0.32, *s.e.* = 0.06), general risk tolerance (*r_g_* = 0.13, *s.e.* = 0.04), and height (*r_g_* = 0.11, *s.e.* = 0.03). The differences in correlations for neuroticism, subjective well-being, and risk tolerance were statistically significant when comparing EA and NonEA-INC, with NonEA-INC showing stronger positive correlations with well-being and risk tolerance and a less negative correlation with neuroticism.

### Implicated genes and tissue-specific enrichments

We used FUMA^42^ to find genes implicated in INC factor GWAS. FUMA uses four mapping approaches: positional, chromatin interaction, expression quantitative trait locus (eQTL) mapping, and MAGMA gene-based association tests. In total, 2,385 protein-coding genes were implicated by at least one of the methods, out of which 225 genes were implicated by all four methods (**Extended Data Fig. 5a**). Only three of these commonly implicated genes were unique for the INC factor, compared to the genes implicated in EA GWAS by at least one of the four methods or previously prioritised for EA.^21^

We then performed tissue-specific enrichment analyses using LDSC-SEG^43^ and MAGMA gene-property analyses^44^ (see **Supplementary Information section 7**). Both methods indicated dominant enrichment for tissues of the central nervous system (**Extended Data Fig. 5b**), consistent with the previous results for household income and EA.^21,24^

## Discussion

We conducted the largest GWAS on income to date, incorporating individual, household, occupational, and parental income measures. Our study design provided increased statistical power, identifying more genetic variants and improving the predictive power of the polygenic index (PGI) compared to previous income GWAS. Additionally, it allowed for comprehensive additional analyses.

We observed substantial heterogeneity in the genetic architecture of income across cohorts and high, but non-perfect genetic correlations of income across sexes. This underlines that the genetic associations we report here are averages across different groups and environments that should not be interpreted as fixed or universal.

Furthermore, we found a strong genetic correlation between income and educational attainment (EA).

Our analyses highlighted numerous associations between better health and higher income that are influenced by genetic differences among individuals. These better health outcomes include lower BMI, blood pressure, type-2 diabetes, depression, and reduced stress-related disorders. We note that the genetic overlap between income and health could be driven by different causal mechanisms, including pleiotropic effects of genes, limited income opportunities for individuals with health problems, or health advantages for individuals with higher income. Investigating these causal mechanisms is outside the scope of this study.

Interestingly, the genetic components of income not shared with EA (NonEA-INC factor) showed weaker associations with better physical health and health-related behaviour, such as drinking and smoking. One possible interpretation of this finding is that better health outcomes of higher socioeconomic status in wealthy countries are more due to their association with education rather than with income or wealth, consistent with findings from quasi-experimental studies.^45–47^

In contrast, we found negative genetic correlations of the NonEA-INC factor with schizophrenia, bipolar disorder, autism, and obsessive-compulsive disorder, while EA exhibited positive genetic correlations with these psychiatric outcomes. This may indicate that the educational system is more accommodating to individuals with these disorders than the labour market and/or that talents associated with these genetic risks (e.g., higher IQ with autism^48^ or creativity with bipolar disorder and schizophrenia^49^) are more advantageous in school than in the labour market.

While our GWAS results contribute to constructing an income-specific PGI with improved predictive accuracy, the EA PGI remains a comparable or even better predictor of income and socio-economic status. This is due to even larger sample sizes in recent GWAS on EA (*N* ∼ 3 million), lower measurement error in educational attainment compared to measures of income and the high genetic correlation between income and EA.

It is important to point out that the results of our study reflect the specific social realities of the analysed samples and are not universal or unchangeable. This is exemplified by the substantial heterogeneity in the genetic architecture of income that we found across cohorts. We emphasise that our results are limited to individuals whose genotypes are genetically most similar to the EUR panel of the 1000 Genomes reference panel compared to people sampled in other parts of the world. Our results have limited generalizability and do not warrant meaningful comparisons across different groups or predictions of income for specific individuals (**FAQ**). To increase the representation of individuals from diverse backgrounds, cohort and longitudinal studies should seek to sample more diverse and representative samples of the global population.

Studies of genetic analyses of behavioural phenotypes have been prone to misinterpretation, such as characterising identified associated variants as ‘genes for income.’ Our study illustrates that such characterisation is incorrect for many reasons: The effect of each individual SNP on income is minimal, capturing less than 0.01% of the overall variance in income. Furthermore, the genetic loci we identified correlate with many other traits, including education and a wide range of health outcomes. Finally, the finding that only one-quarter of the genetic associations we identified are due to direct genetic effects suggests the potential importance of family-specific factors, including potential resemblance between parents, and environmental factors as important drivers of income inequality.

### Box 1. Understanding Genetics and Income: A Cautionary Overview

Given the frequent misunderstanding of research on genetics and human behaviour, it is important to recognize the complexities underlying connections between genes and social outcomes and to communicate what our findings mean clearly and with appropriate nuance.

***What did we do and why?***

Several types of ’luck’ help shape an individual’s life trajectory, such as their society of birth, parents, and the genetic variants they inherit. Our study captures elements of this by examining the relationship between millions of genetic variants and income through a genome-wide association study (GWAS). GWASs of income can provide valuable insights into the genetic factors associated with income and how they interact with environmental factors, enhancing our understanding of intergenerational mobility and socioeconomic disparities.

GWASs of income can shed light on societal processes that favour certain genetic predispositions, providing insights into our socioeconomic system, but also into the relationships between income and health disparities. Recent GWASs have shown that socio-economic outcomes share genetic overlap with various health outcomes, with a considerable portion mediated through social environments.^50^

***What did we find?***

We identified numerous genetic variants associated with income, each with minor effects but collectively correlating with education, cognition, behaviour, and health. We found notable differences between income and educational attainment in their genetic associations with health outcomes. For several psychiatric disorders - namely autism, schizophrenia, and OCD - the genetic relationships acted in opposing directions. Shared genetic effects between income and health may stem from various causes. Genes might affect both income and health. Alternatively, higher income could lead to better health outcomes, not only directly but also indirectly through improved living conditions from family-members or neighbourhoods. Conversely, existing health problems may limit income opportunities, potentially due to reduced work capacity or increased healthcare costs.

When predicting differences between siblings, the overall predictive strength of these genetic effects diminishes significantly — by approximately 75%. Possible explanations for this include that the direct causal effects of the genetic variants are smaller compared to the causal effects of environmental factors that correlate with these genetic variants (e.g., the effects of parental nurture on their children) and that the way parents resemble each other (assortative mating) magnifies the predictive power of genetic effects.

We observed some variability in the genetic factors influencing income across the Western countries we analysed and between genders, underscoring that the genetic associations we report here should not be interpreted as fixed or universal.

### Neither genetic nor environmental determinism is warranted

Historically, misinterpreting the role of genetics in shaping social outcomes has occasionally fueled controversial ideologies with far-reaching consequences. It is important to mitigate the risk of such misunderstandings, particularly the notions of genetic or environmental determinism. In this context, we emphasise the following:

One’s genetic makeup or the family and societal environment into which they are born does not dictate their intrinsic value. The genetic variants that matter for income, and their effects, depend on the environment, i.e., on what skills are valued by the labour market and by society. As the labour market changes, or as government policies change, so can the variants and their effects.

It is important to recognize how genetics can impact income through diverse pathways, affecting one’s own or one’s parents’ health, cognition, skills, and productivity-related behavioural tendencies, such as creativity, risk taking, or adaptability. Additionally, genetics can influence characteristics favoured or discriminated against in the labour market due to societal preferences.

As with previous genetic studies on social outcomes like educational attainment, the findings of this study have limited generalisability across different populations.

## Supporting information

Frequently Asked Questions

## Contribution

H.K., C.A.P.B., T.A.D., R.K.L., A.O., R.D.V., and P.D.K. designed the GWAS meta-analysis. P.D.K oversaw the study. C.A.P.B. was the lead analyst for the meta-analysis, responsible for GWAS, quality control, and meta-analysis. H.K. was the lead analyst for the follow-up analyses, including heterogeneity, MiXeR, GWAS-by-subtraction, genetic correlation, PGI prediction, and biological annotation analyses. N.Y. assisted with the follow-up analyses. R.A conducted PGI prediction and heritability analyses in the STR sample. C.X and W.D.H contributed to several analyses, including cross-country heterogeneity and biological annotation. H.K., D.J.B., and P.K. drafted the manuscript. A.A. wrote Box 1. J.P.B., T.A.D., R.K.L., Q.L., T.T.M., A.O., K.P.H., A.A., W.D.H., and R.D.V. provided important input and feedback on various aspects of the study design and the manuscript. All authors contributed to and critically reviewed the manuscript. The individual contributions of all authors according to the CRediT taxonomy are listed in Supplementary Table 29.

## Methods

This section provides the overall summary of the analysis methods. Further details are available in the Supplementary Information.

### GWAS meta-analysis

We pre-registered our analysis plan for the main income GWAS meta-analysis on August 30 2018 (https://osf.io/rg8sh/). We used four measures of income (individual, occupational, household, and parental income) and conducted a multivariate GWAS to combine these different measures. In total, we recruited 32 cohorts. Some of these cohorts contributed to multiple income measures. **Supplementary Tables 1 and 2** summarise the income measures used for each cohort. **Supplementary Section 2.1** provides details on the phenotype definition. The study was limited to 1KG-EUR-like individuals who were not enrolled in an educational program at the time of survey or who were above the age of 30 if their current enrollment status was unknown.

Each cohort conducted the additive association analysis as follows. The log-transformed income measure was regressed on the count of effect-coded alleles of the given SNP, controlling for any sources of variation in income that do not reflect individual earning potential according to the data availability of each cohort. This included hours worked (with square and cubic terms), year of survey, indicators for employment status (retired, unemployed), self-employment, and pension benefits (see Supplementary Table 4). In addition, the covariates included at least the top 15 genetic principal components and cohort-specific technical covariates related to genotyping (genotyping batches and platforms). This analysis was performed for male and female samples separately.

We applied a stringent QC protocol based on the EasyQC software package^51^ to the GWAS results from each cohort (see **Supplementary Information section 2.4** for more detail). In order to combine multiple GWAS results on different income measures collected from multiple cohorts, we performed the meta-analysis in several steps. First, for each income measure and each sex, we meta-analyzed the cohort-level GWAS results with METAL^27^ using sample-size weighting. Second, for each income measure, we meta-analyzed the male and female results by using the meta-analysis version of MTAG.^28^ To extract the common genetic factor from the four GWAS results with different income measures, we again leveraged MTAG, allowing for different heritabilities among the input traits.

### Cross-sex and cross-country heterogeneity

We investigated the potential environmental heterogeneity in the GWAS of income by estimating the cross-cohort genetic correlations by sex or by country with LDSC.^39^ Sex-specific meta-analysis results for each income measure were available as intermediary outputs from the meta-analysis procedure. In addition, we conducted INC factor GWAS on the sex-specific results, which yielded an effective sample size of 360,197 for men and 353,429 for women.

To derive country-specific GWAS meta-analyses, we only used occupational and household income, for which we were able to obtain a sufficiently large sample size for multiple countries. We obtained the household income GWAS for the USA (*N_eff_* = 30.855), the UK (*N_eff_* = 387,579), and the Netherlands (*N_eff_* = 40,533); and the occupational income GWAS for Estonia (*N_eff_* = 75,682), Norway (*N_eff_* = 42,204), the UK (*N_eff_* = 279,883), and the Netherlands (*N_eff_* = 24,425).

### Comparative analysis with EA

We compared our INC factor GWAS results with the GWAS of EA by examining genetic correlation with LDSC and using the GWAS-by-subtraction approach.^52^ Here, we used a version of EA summary statistics slightly different from publicly available ones. The latest EA GWAS study revised the coding of the years of schooling in the UKB^33^ to better reflect the educational qualification of the participants. Based on the new coding, we conducted a GWAS of EA in the UKB. Then, by using MTAG with the meta-analysis option, we meta-analyzed the UKB result with EA3 summary statistics^21^ that did not include the UKB.

We then statistically decomposed the estimated genetic association of the INC factor into the indirect effect due to EA and the direct effect unexplained by EA (NonEA-INC), leveraging the GWAS-by-subtraction approach in genomic SEM.^29,52^ We implemented this method in the form of a mediation model.

### PGI analysis

We conducted three sets of analyses based on the polygenic index (PGI): 1) prediction analysis, 2) direct genetic effect estimation, and 3) phenome-wide association study of common diseases.

For the PGI prediction analysis, we used the Swedish Twin Registry (STR),^53^ UKB siblings (UKB-sib), and the Health and Retirement Study (HRS).^54^ We constructed PGIs using the meta-analysis results of income excluding a prediction cohort at a time, as well as a PGI based on the EA GWAS summary statistics constructed in the same way for comparison. PGIs were created only with HapMap3 SNPs^55^ as these SNPs have good imputation quality and provide good coverage for 1KG-EUR-like individuals. We derived PGIs based on a Bayesian approach implemented in the software LDpred2.^34^

We measured the prediction accuracy based on incremental *R*^2^, which is the difference between the *R*^2^ from a regression of the phenotype on the PGI and the baseline covariates and the *R*^2^ from a regression on the baseline covariates only. Because income typically contains substantial demographic variation, we pre-residualized the log of income for demographic covariates. Then, as baseline covariates, we only included the top 20 genetic PCs and genotype batch indicators. Because income data was available for multiple years for the STR and the HRS, we residualised the log of income for age, age^2^, age^3^, sex, and interactions between sex and the age terms within each year and obtained the mean of residuals for each individual. For the UKB-sib, which only had cross-sectional data, we residualised the log of income for age, age^2^, age^3^, sex, dummies for survey year, and interactions between sex and the rest. For the EA measure (years of education), we applied the same procedure with birth year dummies. We constructed confidence intervals for the incremental *R*^2^ by bootstrapping the sample 1,000 times.

To estimate the direct genetic effect of the INC factor PGI, we used snipar^38^ to impute missing parental genotypes from sibling and parent-offspring pairs. Parental PGIs were then created with the imputed SNPs. We estimated the direct genetic effect of the PGI by controlling for the parental PGI. This analysis was conducted only with the UKB-sib sample. See Supplementary Information 5.2 for further details.

To explore the clinical relevance of the INC factor PGI for common diseases, we carried out a phenome-wide association study, using the in-patient electronic health records for 115 diseases with sex-specific sample prevalence no lower than 1% in the UKB-sib sample. We derived case-control status according to the phecode scheme by mapping the UKB’s ICD-9/10 records to phecodes v1.2.^56^ We fitted a linear regression of case-control status on the INC factor PGI while controlling for the parental PGIs to capture the direct genetic effects of income PGI. As covariates, we also included the year of birth, its square term, and its interactions with sex, genotype batch dummies, and 20 genetic PCs. Standard errors were clustered by family.

---

**Extended Fig. 1.**
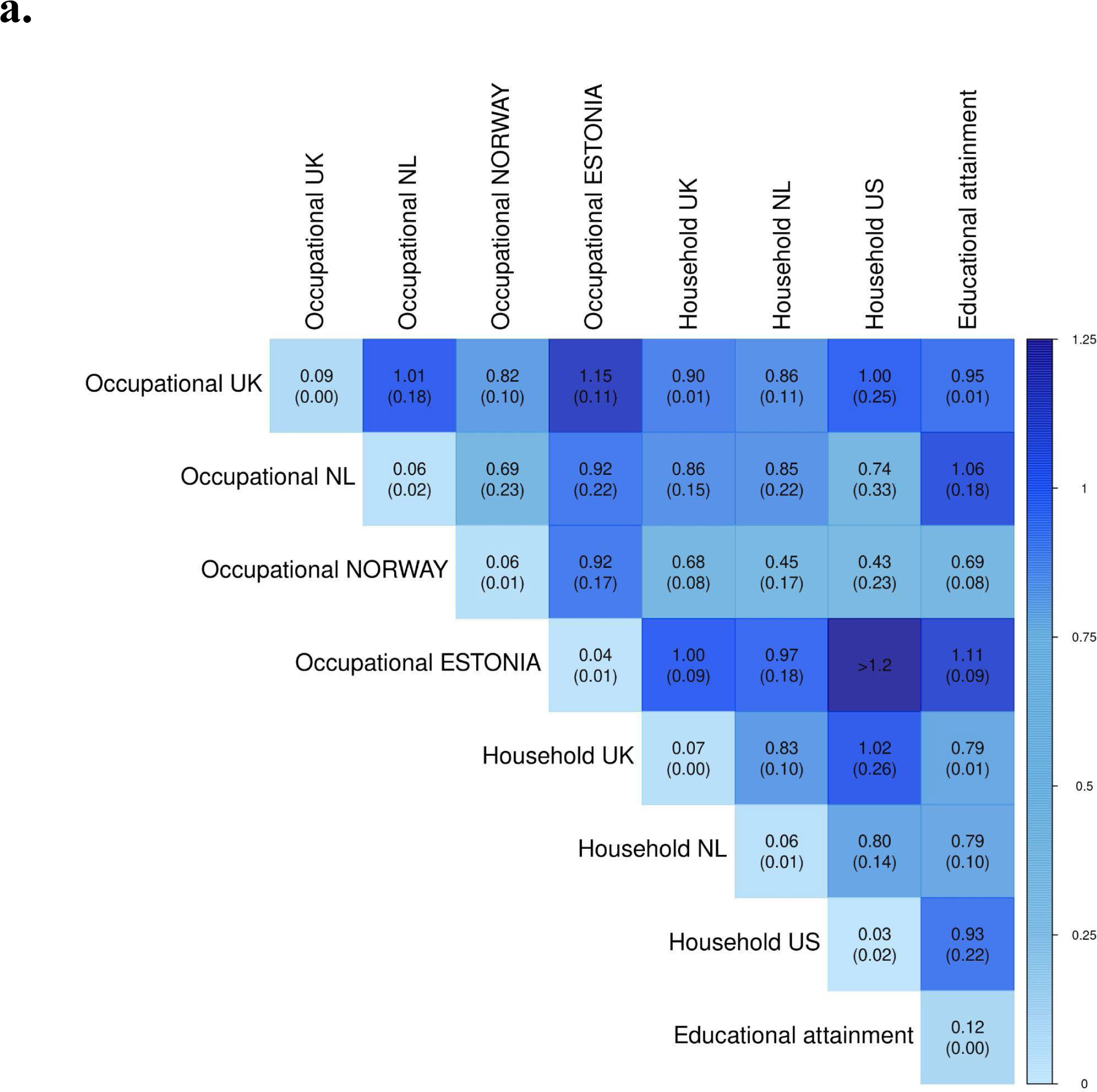
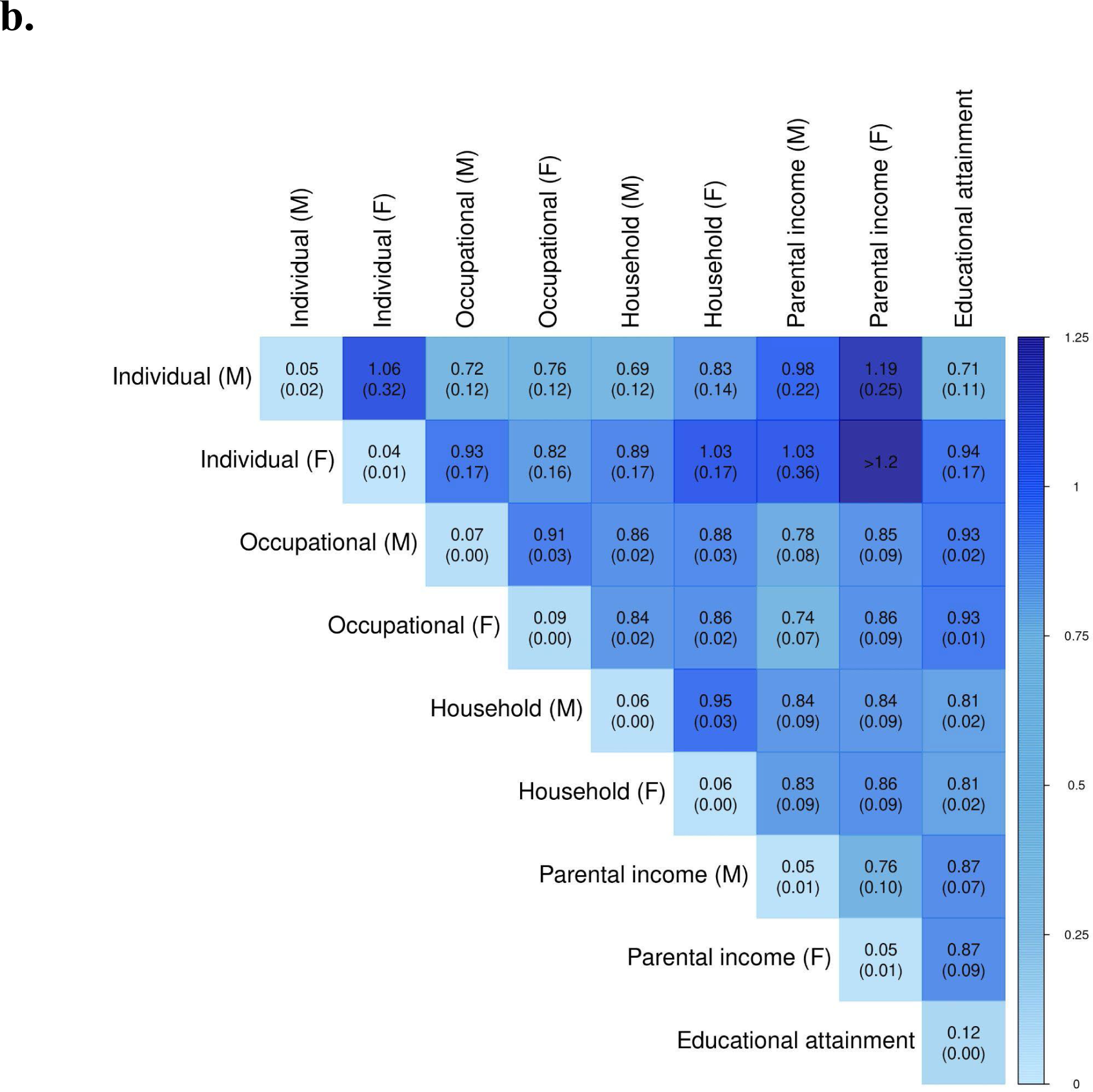
Cross-cohort genetic correlations of income stratified by sex and country. LDSC estimates for cross-cohort genetic correlations of income **a.** between countries and **b.** between male (M) and female (F). The diagonal elements report SNP heritabilities. The standard errors are reported in the parentheses. Some of the results were out-of-bound estimates (exceeding 1.2).

**Extended Fig. 2.**
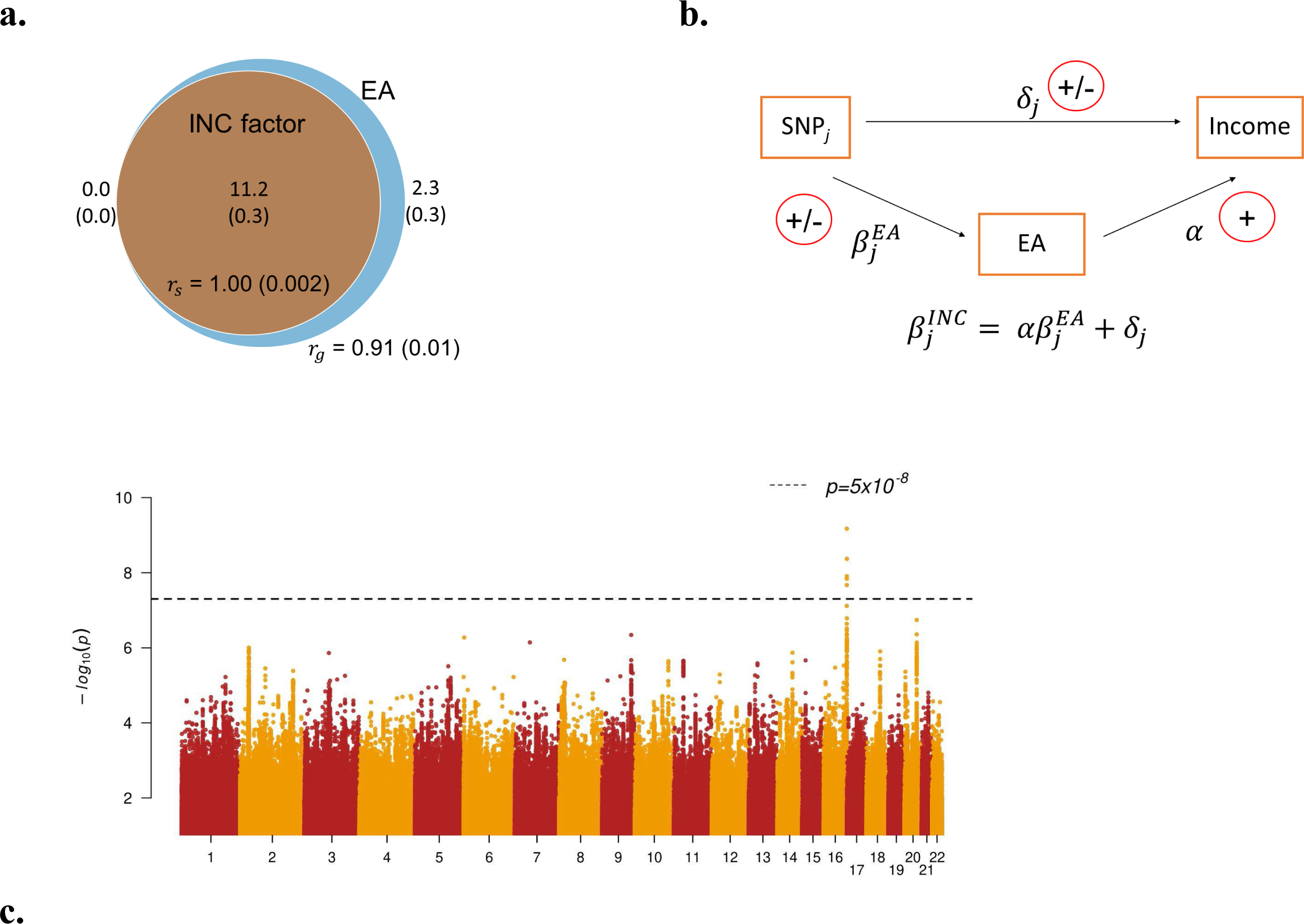
Polygenic overlap of income with EA and GWAS-by-subtraction. **a.** Venn diagram presenting MiXeR results on unique and shared polygenic components for INC factor (orange) and EA (blue). The estimated numbers of unique and shared variants are reported in thousands and by the areas of the circles (0.45 and 2,260 variants for income and EA, respectively; 11,153 shared variants). *r_g_*is the global genetic correlation while *r_s_* is the correlation within the shared variants. The standard errors are reported in the parentheses. **b.** The GWAS-by-subtraction model of non-EA income describes the genetic effect of income for SNP 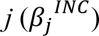 as the sum of two components: 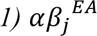: the indirect effect that reflects the genetic association of EA and *2)* δ_j_: the direct effect from SNP to income that reflects the genetic effect of income after statistically removing its genetic covariance with EA. Note that the diagram only depicts a statistical meditation for the sake of interpretation and is not meant to imply any directionality or causal ordering of SNPs to phenotypes. **c.** Manhattan plot showing the non-EA genetic associations of INC factor (NonEA-INC, corresponding to δ_j_ from **b.**). *P* values are plotted on -*log_10_* scale.

**Extended Fig. 3.**
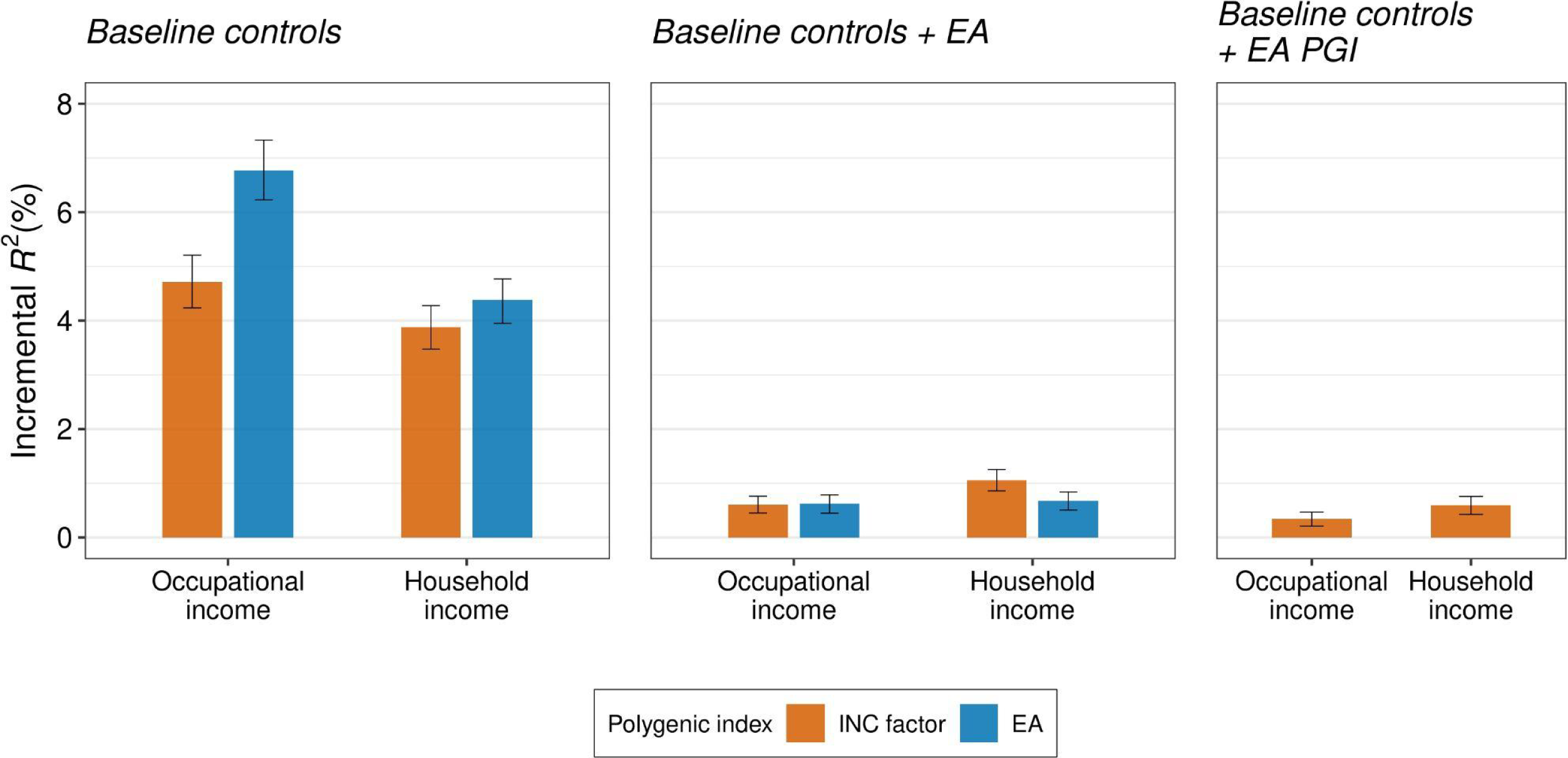
Polygenic prediction of income with additional controls. The figure reports polygenic prediction results in the UKB siblings with PGI for INC factor and additional controls (EA or the PGI for EA). Prior to fitting the regressions, each phenotype was residualized of demographic covariates (a third-degree polynomial in age, year of observation, and interactions with sex). Incremental *R*^2^ is the difference between the *R*^2^ from regressing the residualized outcome on the PGI for INC factor and the controls and the *R*^2^ from a regression only on the controls. The baseline controls include 20 genetic PCs and genotyping batch indicators. Only individuals of European ancestry were included and one sibling from each family was randomly chosen. The error bars indicate 95% confidence intervals obtained by bootstrapping the sample 1,000 times.

**Extended Fig. 4.**
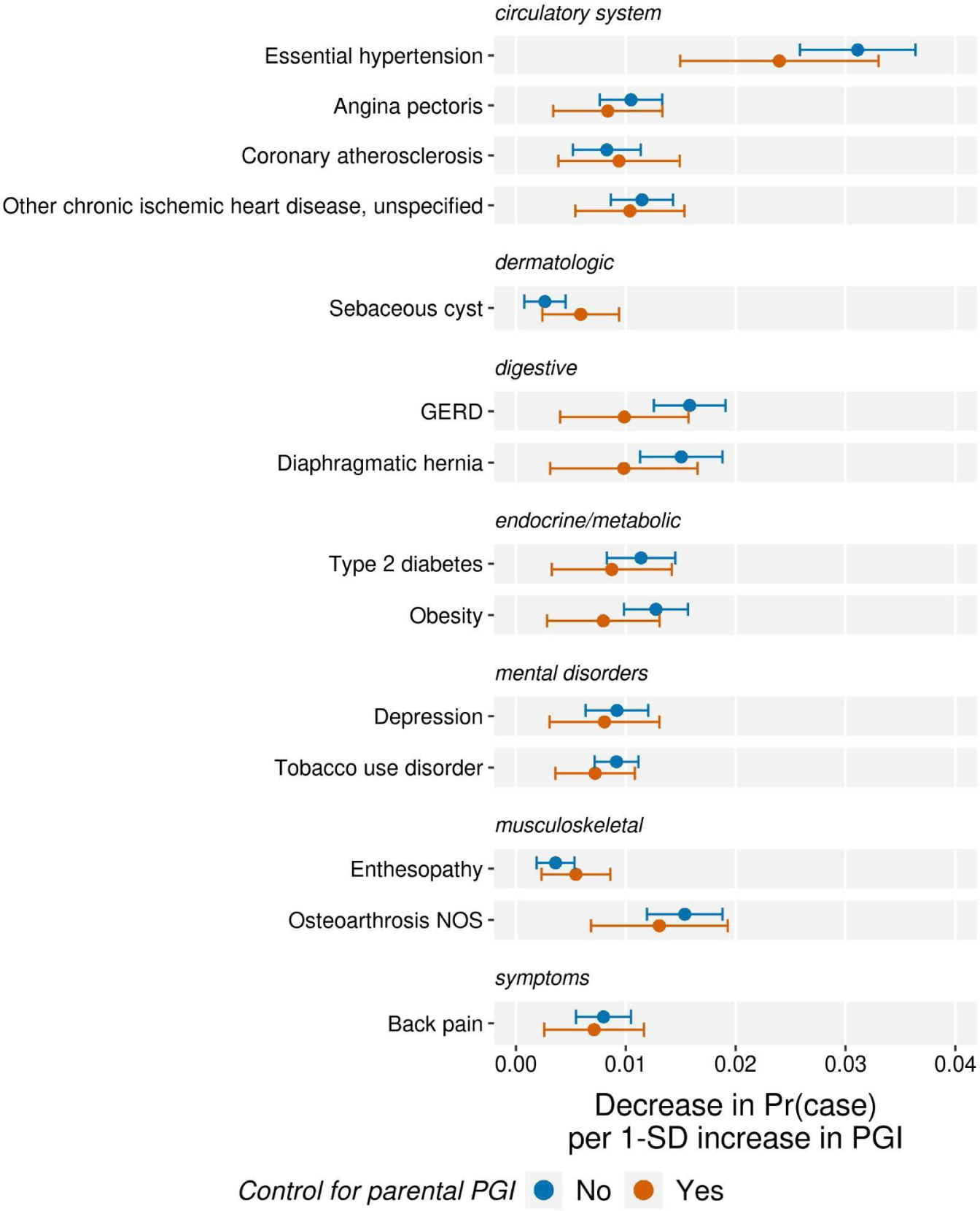
Phenome-wide association study of the INC factor PGI in electronic health records for the UKB sibling sample. The figure reports results from a phenome-wide association study in the in-patient electronic health records of the UKB sibling sample for 115 diseases with sex-specific sample prevalence no lower than 1%. The case-control status was derived according to the phecode scheme by mapping the UKB’s ICD-9/10 records to phecodes v1.2. The case-control status was regressed on the INC factor PGI with and without controlling for the parental PGI. Other covariates included year of birth, its square term, and their interactions with sex, genotype batch dummies, and 20 genetic PCs. The standard errors were clustered by family. The sign of the coefficient estimates was revered to indicate the decrease in the probability of having case status. The results were plotted only for diseases significantly associated with INC factor PGI at the FDR of 5%. with the parental PGI controlled for. The error bars indicate the unadjusted 95% confidence intervals.

**Extended Fig. 5.**
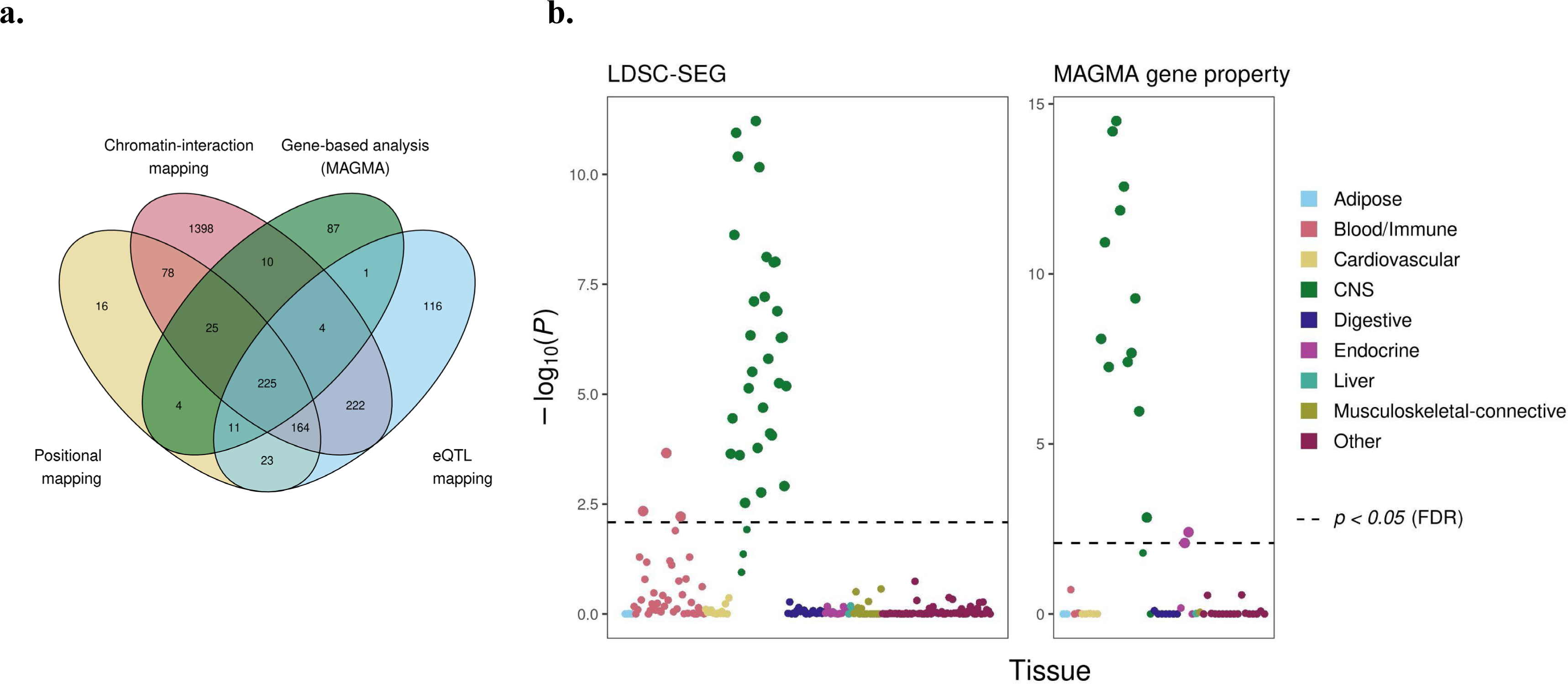
Biological annotation. **a.** The Venn diagram shows the overlap of genes implicated for INC factor by positional mapping, eQTL mapping, chromatin interaction mapping, and MAGMA gene-based analysis. **b.** The figures present the tissue-specific enrichment analysis results based on LDSC-SEG (left) and MAGMA gene-property analysis (right). Each circle indicates a tissue or cell type from either the GTEx or the Franke lab gene expression datasets. Larger circles show statistical significance at the false discovery rate 5%. The full results are reported in Supplementary Table 26.

**Supplementary Fig. 1.**
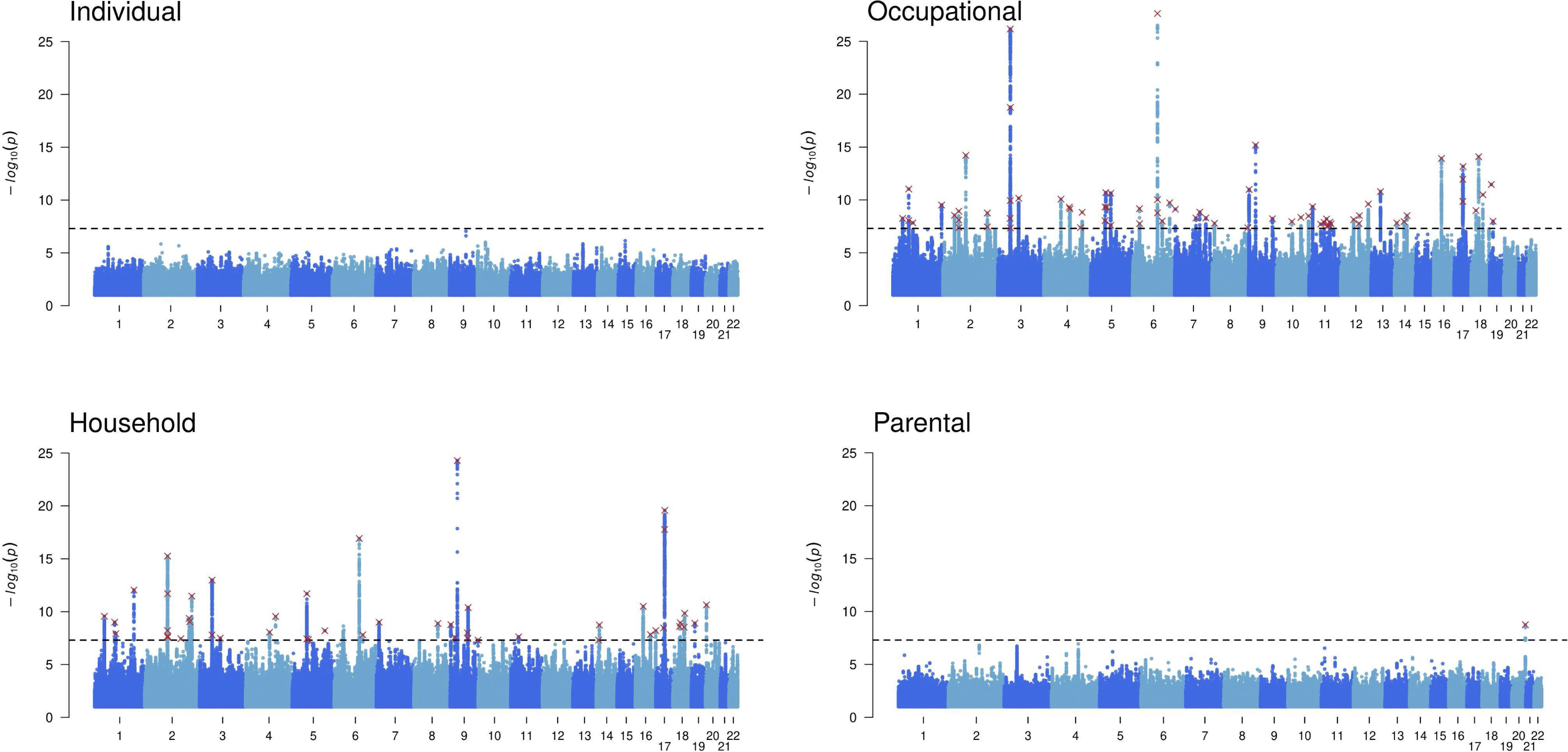
Manhattan plots of income measures.

**Supplementary Fig. 2.**
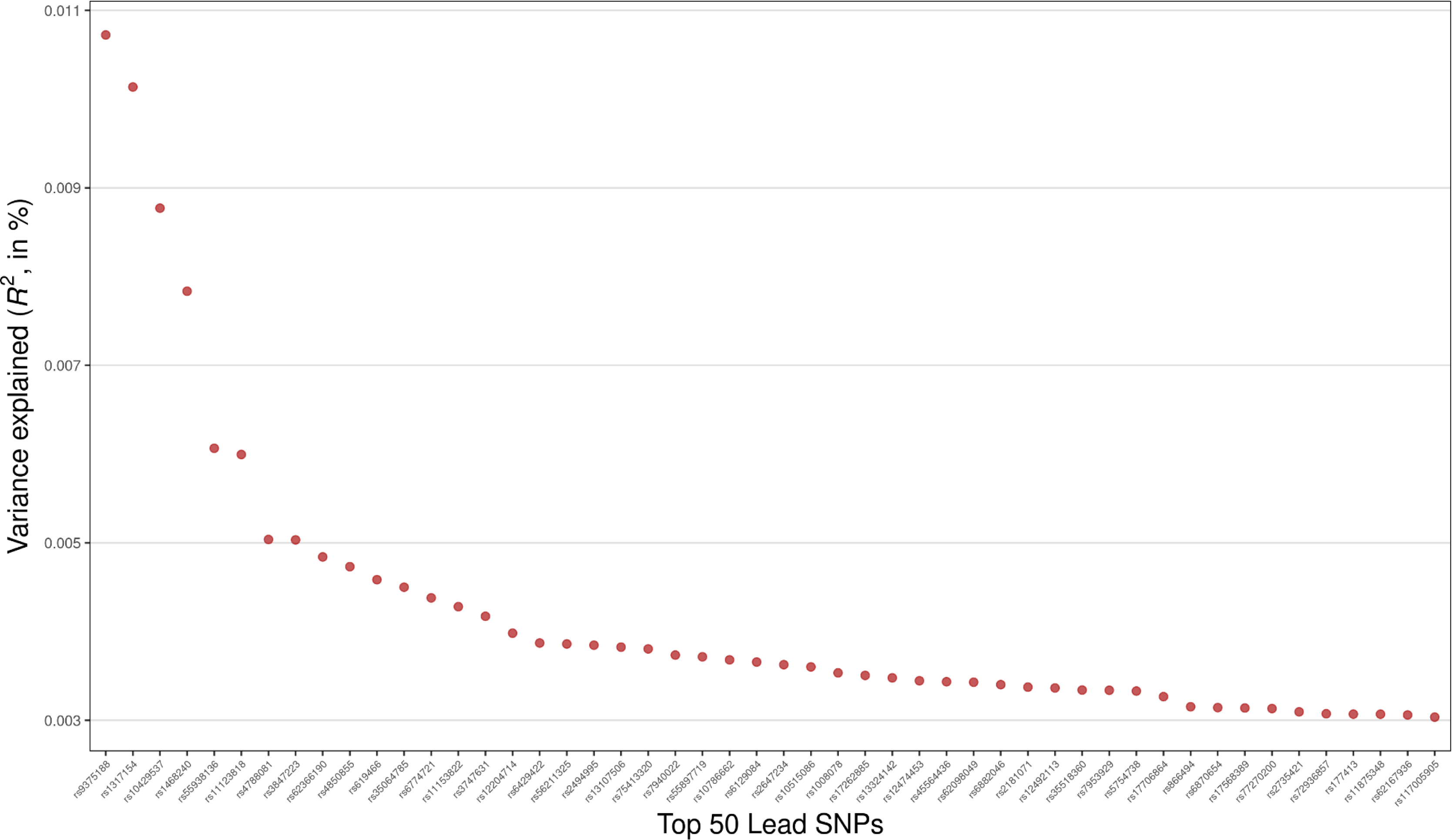
Effect sizes of INC factor GWAS.

