## Supplementary Information for "Associations between common genetic variants and income provide insights about the socioeconomic health gradient"

#### Table of contents

1. Study Overview 2
2. GWAS, quality control, and meta-analysis 3
  2.1. Phenotype definition and construction 3
    2.1.1. General definition 4
    2.1.2. Individual income 4
    2.1.3. Household income 4
    2.1.4. Occupational income 5
      2.1.4.1. UK Biobank and ALSPAC mothers 5
      2.1.4.2. Lifelines and Netherlands Twin Registry 5
      2.1.4.3. Estonian Genome Center 6
      2.1.4.4. HUNT 6
    2.1.5. Parental income (iPSYCH) 7
  2.2. Genotyping and imputation 7
  2.3. Association analyses 7
  2.4. Quality control 7
  2.5. Meta-analysis 8
  2.6. Identification of genomic loci 9
  2.7. Winner's-curse adjustment 10
3. Environmental heterogeneity 10
  3.1. Between-sex heterogeneity 10
  3.2. Cross-country heterogeneity 11
4. Comparison with educational attainment 11
  4.1. LDSC and MiXeR 11
  4.2. GWAS-by-subtraction 12
  4.3. Concordant and discordant sets 13
5. Polygenic score analyses of income 14
  5.1. Baseline polygenic prediction 14
    5.1.1. The STR income data description 15
  5.2. Within-family polygenic prediction 15
6. Genetic correlation analysis 16
7. Biological annotation 17
  7.1. Gene mapping 17

7.2. Tissue-specific enrichment analysis 17
8. GREML heritability estimation 17
9. Phenome-wide association study 17
10. Cohort acknowledgements 18
10.1. ALSPAC (Avon Longitudinal Study of Parents and Children) 18
10.2. CoLaus (Cohorte Lausannoise) 19
10.3. Croatia - Korcula 19
10.4. EGCUT (Estonian Genome Center, University of Tartu) 19
10.5. FTC (Finnish Twin Cohort) 20
10.6. HUNT (Trøndelag Health Study) 20
10.7. iPSYCH 20
10.8. LifeLines 20
10.9. MOBA (Norwegian Mother, Father and Child Cohort Study) 21
10.10. NEO (The Netherlands Epidemiology of Obesity Study) 21
10.11. NTR (Netherlands Twin Registry) 22
10.12. QIMR (Queensland Institute of Medical Research) 22
10.13. RS (Rotterdam Study) 22
10.14. SHIP (Study of Health in Pomerania) 22
10.15. STR (Swedish Twin Registry) 23
10.16. UKHLS (Understanding Society) 23
11. References 23

### 1. Study Overview

We conducted a genome-wide association study (GWAS) of income, using four income measures and data collected from more than 600,000 participants in 26 cohorts from 12 countries. Due to data availability and statistical power considerations, our analyses were restricted to individuals carrying genotypes most similar to the EUR panel of the 1000 Genomes data set (1KG-EUR), as compared to individuals samples elsewhere in the world. The meta analysis across cohort-level results was carried out per each income measure and then the results across the four income measures were combined by extracting their shared genetic basis. Then a number of follow-up analyses were conducted. First, we investigated the environmental heterogeneity between sex and between countries. Second, we examined the shared genetic factor with educational attainment (EA). Third, we performed polygenic prediction analyses. Fourth, we estimated the genetic correlation of income with a number of other related phenotypes and compared the results with the estimates for EA. Finally, we performed brief biological annotation analyses.

This study was carried out under the auspices of the Social Science Genetic Association Consortium (https://www.thessgac.org/).

#### 2. GWAS, quality control, and meta-analysis

We pre-registered our analysis plan for the main income GWAS meta-analysis on August 30 2018 (https://osf.io/rg8sh/). In total, we recruited 31 cohorts, which have one of the following income measures available: individual, occupational household, and parental income. Some of these cohorts contributed to multiple income measures.

The sample inclusion criteria according to our analysis plan are as follows:

1. Samples are of European ancestry (1KG-EUR-like individuals);
2. They are finished with education. If such information is unavailable, limit analyses to those aged >30 years
3. All relevant covariates are available for the individual
4. They were successfully genotyped genome-wide (recommended individual genotyping rate: > 95%).
5. They passed the cohort-specific standard quality controls (e.g., excluding individuals who are genetic outliers in the cohort).
6. In the case of self-reported income, unreasonable answers should be removed (e.g., negative income or yearly income > 10 mio EUR). The number of deleted observations and the respective reason for deletion as well as income histograms need to be reported in the descriptive statistics summary file of the cohort.

##### 2.1. Phenotype definition and construction

Individual income is the result of various factors including achieved qualifications (e.g. education, learnt occupation, experience), personal characteristics (e.g. leadership, cognitive skills, consciousness), the demand and supply for these qualifications and characteristics in the labor market, and personal choices about labor supply (e.g. due to personal preferences, decisions about division of labor among household members). In this paper, we aimed to study the genetic factor for such individual earning potential. For this purpose, it was ideal to use individual income measures. However, individual income information was typically not collected in most of the genotyped samples. To circumvent such empirical challenges, we used four measures of income (individual, occupational, household, and parental income) and conducted a multivariate GWAS to combine these different measures. **Supplementary Tables 1-2** summarize the details of income measures used for each cohort.

###### 2.1.1. General definition

For all income measures considered, we defined the main phenotype as the natural log of income before-tax. It is important to use the log transformation here because this allows us to correct for the typical skewness of the income distribution, which will return a better linear fit, as well as to model the percentage change in income, which is unit-free. Ideally, the phenotype included all “earned" financial compensation (salaries, income from self-employment, profits from running one’s own business, bonuses, vacation benefits) but excluded non-earned monetary transfers such as rental income, capital gains, dividends, and transfers from the government, family, or former spouses.

Many cohorts opted to use categorical responses to measure individual or household income. In these cases, we converted these categories to a semi-continuous measure by taking the natural logarithm of the midpoint of the category. As the top and bottom category are often open-ended and do not have a midpoint, we converted the top category by taking the logarithm of 4/3 times the lower bound of that category and the bottom category by taking the logarithm of 3/4 times the upper bound of that category.

When multiple observations of the income measure per individual were available (i.e. longitudinal data), we first regressed the income measure on all control variables including time-specific intercepts. Then, the mean of the residuals for each person were taken as the phenotype.

Some of the cohorts of older adults had a large share of retired individuals who may have been receiving pension. For these individuals, we used their last observed wage. If their last wage was not available, we derived occupational wage from their last occupation. In either case, they were treated as if they were observed while they had their last job. For instance, if a 65-year-old retired individual was surveyed in 2009 and her past wage or occupational wage for the job that she had in 2006 was available, her age and year of observation was 62 and 2006, respectively, in the control variables.

Individuals who are unemployed or economically inactive at the time of survey were treated like pensioners if they had an income in the past. In other words, their last observed income or occupation was used.

###### 2.1.2. Individual income

Official registry data (e.g. from tax records) are most ideal to obtain high-accuracy measures of individual income. However, the linkage between genetic data and registry data was normally not feasible due to privacy concerns. Therefore, we mainly relied on self-reports of income, despite likely measurement error.

###### 2.1.3. Household income

We considered household income as an alternative measure of individual income. Household income aggregates the individual incomes of all household members (e.g. spouses and possibly even children or other relatives). Therefore, household income captures not only factors that contribute towards individual income, but also other factors such as the ability and desire to attract a spouse and the characteristics of the spouse. Nonetheless, household income can still serve as a reliable proxy of individual income.

##### 2.1.4. Occupational income

When detailed occupation information was available with standardized coding, we derived (the logarithm of) occupational income based on the national statistics data for each country. Occupation encompasses income potential and typically also reflects educational attainment, personal interests, social prestige and labor market opportunities. In comparison to individual income, occupational income only captures between-occupation variation in individual income. However, occupational income is less likely to suffer large measurement error because it is easier to recall occupation than income, while occupation-specific income is obtained from the national statistics of the relevant country. Occupational income measures were mainly used for larger cohorts. Due to different data availability across different countries in which those cohorts are based, slightly different approaches were used for different cohorts, which are summarized below.

###### *2.1.4.1. UK Biobank and ALSPAC mothers*

The UK Biobank recorded the occupation of participants with the UK's standardized occupational classification (SOC) 2000 version, which is coded in 4-digit numbers representing a hierarchical structure. Similarly, ALSPAC also provided occupational information in the same coding for the mother participants, while their income was not surveyed. For these British cohorts, we applied the approach that we developed in ref[1]. This approach was originally developed to impute income based on occupation and demographic information, rather than to derive occupational wage. The income imputed this way can be interpreted as expected income per occupation adjusted for demographics, which therefore is not essentially different from occupational income.

The details of the approach are available in the appendix of ref[1]. Here we only provide the overall summary. From the Annual Survey of Hours and Earnings, we obtained the tax-registry-based estimates of sex-specific mean and median hourly wages for each occupational group defined by 4-digit level SOC. Using the Labour Force Survey (LFS), a large representative survey data of the UK population, we fit a regression model of log hourly wages using mean and median wages for each occupation along with demographic variables and interaction terms. The log occupational wages were then derived as the predicted outcomes from this regression. In the appendix of ref[1], it was shown that occupational wages constructed from this method yielded an out-of-sample $R^2$ = 0.50 with self-reported log hourly wages in British Household Panel Survey, another independent representative survey of the UK.

###### *2.1.4.2. Lifelines and Netherlands Twin Registry*

A similar approach was taken for two Dutch cohorts: Lifelines and Netherlands Twin Registry (NTR). We mirrored the approach for the British cohorts as closely as possible. Here we used data from the Dutch Labour Force Survey, 'Enquête Beroepsbevolking' (EBB). The EBB is a national representative survey of the Dutch labor force, conducted by Statistics Netherlands (CBS). We used a merged dataset containing 479,893 individuals in yearly waves from 2012 to 2017, where we excluded multiple observations per individual by taking the latest observation. The EBB used a Dutch version of standardized occupation codes, BRC, developed by CBS based on the International Standardized Classification of Occupation (ISCO) 08 standard.

As the EBB was the only national representative survey containing standardized occupation codes, we fitted a regression model and calculated the mean and median hourly wages per occupation group in the same sample. We standardized hourly wages to the year 2012 using the consumer price index calculated by CBS. We then calculated the mean and median wage for each 4-digit occupation code separately for each sex. If there are less than 10 people per occupation code, we calculated the mean and median using a pooled sample of both sexes. If there are less than 10 people per occupation code in the pooled sample, we used the 3-digit occupation code instead. If the 3 digit occupation code still did not yield a sufficient sample size, we used the 2-digit occupation code. The same model specification as the UK model was used for the wage prediction model.

Given the estimated model, we constructed the log hourly wages per occupation in the NTR and LifeLines. The accuracy of the model was tested by taking the 2017 EBB subset as a hold-out sample ($N$ = 91,821) and re-estimating the regression model using the 2012 – 2016 subset excluding those present in the 2017 ($N$ = 388,072). Regressing the log hourly wage on the imputed log hourly wage in the 2017 EBB subset yielded an $R^2$ of 0.47, which is similar to that for the UK case above.

*2.1.4.3. Estonian Genome Center*

For the Estonian Genome Center (EGCUT), we employed a simpler algorithm. We used the mean log wage of each occupation code, estimated for men and women separately, using the 2011 population census data from Statistics Estonia. EGCUT used 3-digit occupation codes based on the ISCO-88 standard while Statistics Estonia used occupation codes based on the ISCO-08 standard. The mean log wages for each ISCO-08 code were matched to the ISCO-88 codes based on the correspondence file published by the International Labour Organisation. When multiple ISCO-08 codes corresponded to a single ISCO-88 code, we took the average of the estimated means of the ISCO-08 codes.

We tested the accuracy of the occupational wage estimates by examining their correlation with the self-reported log wages in the Structure of Earnings Survey ($N$=369,247 individuals aged 25 to 64). This resulted in $R^2$ = 0.44, which is similar to the results of the Dutch and British cases.

*2.1.4.4. HUNT*

For the Norwegian cohort HUNT, we used a similar approach to that for EGCUT. Here, we used sex-specific mean wage statistics from 2015 to 2019 from the Statistics Norway (https://www.ssb.no/en/statbank/table/11418/). Similarly to the case of EGCUT, HUNT used 3-digit occupation codes based on the ISCO-88 standard while Statistics Norway used occupation codes based on the ISCO-08. The two are matched together in the same way as was done for EGCUT.

##### 2.1.5. Parental income (iPSYCH)

While the income information of the participants of iPSYCH was available, they were too young that their current income was unlikely to reflect their life-time earnings potential. Therefore, we opted to use the income of their parents instead, which was collected from the Danish registry data. Specifically, we used the average earnings of the age 30 ~ 55 for each parent. This approach can be interpreted as using the offspring genotype as a proxy for the genotype of the parent.

#### 2.2. Genotyping and imputation

**Supplementary Table 3** reports cohort-level information on the genotyping platform, quality-control filters for the genotype data and subjects prior to imputation, subject-level exclusion criteria, and the reference panel and software used for imputation. As the reference panel for imputation, either the 1000 Genomes Project[2] or Haplotype Reference Consortium (HRC)[3] was used except for a few cohorts that additionally used cohort-specific reference data.

#### 2.3. Association analyses

Each cohort estimated the following linear regression model for each SNP.

$$y_i = \beta_0 + \beta_1 SNP_i{}^j + Z_i{}'\gamma + \varepsilon_i$$

$y_i$ is the log-transformed income phenotype for individual $i$, $SNP_i{}^j$ the count of effect-coded allele of the SNP $j$, $Z_i$ the vector that contains control variables with corresponding coefficients $\gamma$, and $\varepsilon_i$ the error component. Each cohort was asked to control for any sources of variation in income that do not reflect individual earning potential according to their data availability. This includes hours worked (with square and cubic terms), year of survey, indicators for employment status (retired, unemployed), self-employment, pension benefit, and etc (see Supplementary Table 4). Importantly, each cohort was asked to include at least top 15 genetic principal components (PC) to account for population stratification, as well as cohort-specific technical covariates related to genotyping (genotyping batches and platforms). For household income, the number of adult members was also controlled for if possible.

This model was estimated for male and female samples separately in light of the possible between-sex heterogeneity. Generally, the linear mixed model approach was preferred, which additionally models the error component with random genetic effects in order to account for the family structure and cryptic relatedness. The cohorts were advised to use BOLT-LMM[4] for implementation. For smaller family-based cohorts, for which BOLT-LMM's approximation approach was not expected to work well, fastGWA[5] was used instead. Otherwise, the association analysis was performed without the random effect component.

#### 2.4. Quality control

We applied a stringent quality-control (QC) protocol to each set of GWAS results of each cohort based on the EasyQC software package (version 9.2) developed by the GIANT consortium[6], as well as additional steps developed by the SSGAC[7–9]. As the reference panel, we used HRC v.1.1[3]. All issues raised during the QC protocol were resolved through iterations with cohort analysts, before the meta-analyses.

The details of the QC protocols as well as the QC of the HRC reference panel is described in the supplementary materials of ref[9]. Here we only provide the overall summary. The main steps include removing SNPs with missing or incorrect numerical values (a *p*-value outside of [0,1], for instance); a minor allele frequency (MAF) below 0.1% or a minor-allele count (MAC) below 200; a low imputation accuracy (0.6 for MACH, 0.7 for IMPUTE, 0.8 for PLINK); the effect-coded allele or the other allele with values different from "A," "C," "G," or "T."; a Hardy-Weinberg Equilibrium *p*-value lower than $10^{-3}$ ($N$<1000), $10^{-4}$ (1000≤ $N$<2000), or $10^{-5}$ (2000≤ $N$<10000) ; and an allele frequency different from the allele frequency in the reference panel by more than 0.2. We also removed duplicate SNPs or SNPs absent in the reference panel.

After applying these steps, the resulting output was inspected to determine if an unusual number of SNPs were removed during one of the steps and when necessary errors were resolved together with the cohort analysts.

#### 2.5. Meta-analysis

In order to obtain a single GWAS output that combines multiple GWAS results on different income measures collected from multiple cohorts, we performed the meta-analysis in several steps, as follows.

Step 1. For each income measure and for each sex, we meta-analyzed the cohort-level GWAS results with METAL using sample-size weighting, which resulted in 8 sets of GWAS summary statistics given the four income measures.

Step 2. For each income measure, we meta-analyzed the male and female results by using the meta-analysis version of MTAG, which specifies the perfect genetic correlation and equal heritability among the input traits. This version of MTAG can be interpreted as a generalized inverse-variance-weighted meta-analysis. In addition to the variance of the estimates, MTAG exploits additional information from the intercepts of LD score regressions to compute the weights and standard errors. This approach helped account for the unadjusted relatedness between the male and female samples. Prior to running MTAG, we dropped the SNPs with $N = N_{male} + N_{female}$ smaller than 50% of the maximum $N$ to make sure that there were no SNPs with an excessively smaller sample size.

Step 3. To combine the four GWAS results with different income measures, we again leveraged MTAG with the perfect genetic correlation specification while allowing for different heritability among the input traits. This approach allowed us to meta-analyze results from different measures that may have different heritability or measurement error as well as to account for the sample overlap, which was important given that some individuals were included in multiple GWAS results with different income measures.

As opposed to the meta-analysis with METAL, MTAG, a multivariate analysis tool, can only output the common set of SNPs among the input GWAS summary statistics. This led to a considerably low number of SNPs (4,885,528) after Step 3 due to the individual income and parental income GWAS results, which did not have any biobank-scale cohort and therefore had a smaller coverage over the genome.

To circumvent this issue, we repeated Step 3 without 1) individual income, 2) parental income, and 3) both individual and parental income. We first verified that all of the four sets of meta-analyzed results, including the one with all the measures, had pairwise genetic correlation estimates larger than 0.99 and their heritability estimates were almost identical from LDSC[10]. These results indicate that the multivariate meta-analysis results are not sensitive to dropping individual and/or parental income. Therefore, for each available SNP, we took the result that gave the largest *Z* statistic in absolute value among the four results. As a result, we obtained 4,885,528 SNPs from the MTAG result with the all four measures and 6,599,628 SNPs from the MTAG result which only includes occupational and household income. We dropped 2,353,649 SNPs whose effective sample size (see below) fell below 70% of the maximum effective sample size (=692,936). In total, 9,131,507 SNPs were included in the final output.

We observed that MTAG with the perfect genetic correlation specification yields numerically almost equivalent results with genomic SEM's default common factor function[11]. We applied genomic SEM's common factor function to extract a common underlying factor of the four income measures. Using a set of SNPs established by the International HapMap 3 consortium[12], we found that the *Z* statistics from the meta-analysis result from MTAG (Step 3) had an $R^2$ of 0.998 with the *Z* statistics from the genomic SEM's common factor results. The mean $\chi^2$ statistics was slightly higher for MTAG (2.118 versus 2.108). In light of this result, we refer to the meta-analyzed income as 'the income factor' (INC factor) to emphasize that this result reflects the shared genetic factor underlying multiple income measures.

We computed the effective sample size exploiting the fact that the standardized beta estimates can be approximated as $Z/\sqrt{N}$ for large *N*. Using the MTAG-produced standardized estimates $\beta_{std}$, we computed the effective sample size per SNP as follows:

$$N_{eff} = (\frac{Z}{\beta_{std}})^2$$

In the downstream analyses, we used these per-SNP effective sample sizes since typical GWAS softwares re-compute the standardized estimates from the MAF, *N*, and *Z* statistic based on the same approximation. To evaluate the overall sample size, we took the average of these per-SNP effective sample sizes using the SNPs with $0.1 < MAF < 0.4$ since these SNPs tend to be less noisy. As a result, we estimated the overall sample size of our INC factor GWAS to be 668,288.

Since MTAG already applies a bias-correction with the intercept of LD score regression, we did not apply further bias adjustments. Also, to measure the effect sizes, we used the (partial) coefficient of determination ($R^2$), which is the square of the standardized beta estimates.

#### 2.6. Identification of genomic loci

We used FUMA v1.5.2[13] with the default parameters to define genomic loci associated with INC factor. Here, we briefly explain the procedure. See the original paper for more detail. FUMA first finds independent significant SNPs with a genome-wide significance ($p < 5x10^{-8}$)

such that they are independent from each other at $r^2 < 0.6$. Then, independent lead SNPs are identified from the independent significant SNPs such that they are independent from each other at $r^2 < 0.1$. FUMA then forms genomic loci by grouping independent significant SNPs if they are less apart than 250 kb. As a result, a genomic risk locus can contain multiple independent significant SNPs and multiple lead SNPs. To define the border of each locus, SNPs that have $r^2 \geq 0.6$ are identified for each of the independent significant SNPs from the reference data, including those not available in the input GWAS summary statistics. We used the 1000 Genomes Project as the reference data, which is readily available in FUMA.

##### 2.7. Winner's-curse adjustment

We used an empirical Bayes framework to adjust for winner's curse bias in the estimated effect sizes from the lead SNPs, following the approach described in ref[7,14]. The marginal effect sizes of SNPs are assumed to be drawn from the following mixture distribution: $N(0, \tau^2)$ with probability $\pi$ and 0 otherwise. Here $\pi$ is the fraction of the non-null SNPs and $\tau^2$ is their effect-size variance.

We estimated the parameters $\pi$ and $\tau^2$ by maximum likelihood, using all the SNPs from our INC factor meta analysis, which yielded $\hat{\pi} = 0.65$ and $\hat{\tau}^2 = 3.01 \times 10^{-6}$. On average among the SNPs with $0.1 < MAF < 0.4$, this corresponds to the shrinking factor of 0.67, which implies that we need to shrink the GWAS effect estimate by 33% to obtain the winner's-curse-adjusted estimate, conditioning that the SNP is non-null. For full technical details of these derivations, see the Supplementary Note of ref[7,14].

We then computed the 5th, 50th and 95th percentile of the effect-size distribution of the lead SNPs as follows. We simulated 10,000 effect sizes from the posterior distribution of each lead SNP and obtained the 5th, 50th and 95th percentiles from the complete set of simulated effect sizes.

#### 3. Environmental heterogeneity

We investigated the potential environmental heterogeneity in the GWAS of income by examining the cross-cohort genetic correlations by sex or by country.

##### 3.1. Between-sex heterogeneity

We estimated genetic correlation using LDSC between male and female meta-analysis results for each income measure (from Step 1 in Section 2.5). In addition, we conducted INC factor GWAS on the sex-specific results (Step 3 in Section 2.5), which yielded an effective sample size of 360,196.7 for men and 353,429.1 for women. We then estimated the genetic correlation between the sex-specific INC factor results.

##### 3.2. Cross-country heterogeneity

We derived country specific GWAS meta-analyses on two measures of income, occupational and household income, for which we were able to secure a sufficiently large sample size for multiple countries. We applied Step 1 and 2 in Section 2.5 using the cohorts from each country. As a result, we obtained the household income GWAS for the USA ($N_{eff}$ = 30.855), the UK ($N_{eff}$ = 387,579), and the Netherlands ($N_{eff}$ = 40,533); and the occupational income GWAS for Estonia ($N_{eff}$ = 75,682), Norway ($N_{eff}$ = 42,204), the UK ($N_{eff}$ = 279,883), and the Netherlands ($N_{eff}$ = 24,425). We then estimated pairwise gene`tic correlations between these results with LDSC.

Next, we examined whether our meta-analysis results could be driven by the cohorts from the UK due to a dominantly large share of the British cohorts in the meta-analysis. We repeated the meta-analysis procedure (Step 1-3 in Section 2.5) separately with the British and non-British cohorts (from 11 countries), which yielded two GWAS results for INC factor. The effective sample size was 414,978 for the UK result and 330,639 for the non-UK result. We then estimated the genetic correlation between them.

#### 4. Comparison with educational attainment

We compared our INC factor GWAS results with the GWAS of educational attainment (EA, measured as years of education) in several approaches by examining 1) genetic correlation with LDSC, 2) polygenic overlap with MiXeR[15], and 3) GWAS-by-subtraction[16].

Here, we used a version of EA summary statistics that is slightly different from those publicly available. The latest EA GWAS study[17] revised the coding of the years of schooling in the UKB, which better reflects the educational qualification of the participants. We conducted a GWAS of EA in the UKB based on the new coding. Then, by using MTAG with the meta-analysis option, we meta-analyzed the UKB result with EA3 summary statistics that did not include the UKB. This increased the mean $\chi^2$ from 2.53 of the original EA3 result to 2.94. We found 872 loci tagged by 1,473 lead SNPs.

##### 4.1. LDSC and MiXeR

Using LDSC[10], we estimated that the genetic correlation between INC factor and EA is 0.92 (*s.e.* = 0.01). This result was consistent with the previous reports, which ranged from 0.90 to 0.94[1,18,19]. Though providing a useful summary of the shared genetic basis, the global genetic correlation only estimates the average correlation of genetic associations and does not capture mixtures of effect directions.

To gain further insights, we used MiXeR[15] tool to estimate the degree of polygenic overlap between INC factor and EA. MiXeR exploits a bivariate causal mixture model, which allows for estimating: 1) the number of non-null SNPs specific to each trait, 2) the number of SNPs with non-null effect for both traits, and 3) the genetic correlation within the shared variants.

More specifically, MiXeR models GWAS effects as a mixture of four components: *1)* SNPs with null effects for both traits, *2)* non-null SNPs specific to trait 1, *3)* non-null SNPs specific to trait 2, and *4)* SNPs with non-null effects for both traits. Each of the four components is represented by the proportion of its member SNPs denoted by $\pi_o$, $\pi_1$, $\pi_2$, and $\pi_{12}$, respectively. The SNPs in the second and third components (non-null SNPs specific to each trait) are assumed to be distributed with $N(0, \sigma_k^2)$ for trait $k$ = 1 and 2, respectively. The SNPs in the fourth component (the shared non-null SNPs) are distributed with a bivariate normal distribution with the variance-covariance matrix:

$$\begin{bmatrix} \sigma_1^2 & \rho_{12}\sigma_1^2\sigma_1^2 \\ \rho_{12}\sigma_1^2\sigma_1^2 & \sigma_2^2 \end{bmatrix}$$

where $\rho_{12}$ indicates the correlation of the GWAS effects within the shared SNPs.

Under the MiXeR model, the global genetic correlation is estimated as: $\rho_{12}\pi_{12} / \sqrt{(\pi_1 + \pi_{12})(\pi_2 + \pi_{12})}$, which is the correlation of effects within the shared variants scaled by the normalized degree of their polygenic overlap. Therefore, a low global genetic correlation may indicate a low correlation within the shared variants and/or a low degree of polygenic overlap.

The estimated model suggests that the set of SNPs associated with the INC factor is entirely nested within the set of SNPs associated with EA, with 83.2% of the EA SNPs shared with income (**Extended Data Fig. 2a**). Furthermore, the genetic correlation within the shared component is perfect ($r_g$ = 1.00, s.e. = 0.002). As a result, the global genetic correlation, which is a composite measure of the polygenic overlap and genetic correlation within the shared set, was estimated to be 0.91 (s.e. = 0.01), consistent with the LDSC result.

#### 4.2. GWAS-by-subtraction

To explore their results from MiXeR, we statistically decomposed the estimated genetic association of INC factor into the indirect effect due to EA and the direct effect unexplained by EA (denoted 'NonEA-INC' hereafter), using the GWAS-by-subtraction approach[16]. While this method was implemented as a Cholesky model in the original study, we implemented this in a form of the mediation model, which produces numerically equivalent results. The main difference is that we did not specify latent factors whose variance was fixed to unity. This therefore only affects the scale of the beta estimates and standard errors, while the Z statistics are the same.

More specifically, we set up a genetic mediation model of INC factor with EA as an mediator (**Extended Data Fig. 2b**). Under this model, the genetic association of INC factor for SNP $j$ ($\beta_j^{INC}$) can be written as $\beta_j^{INC} = \alpha \times \beta_j^{EA} + \delta_j$, where each component is defined as:

- $\alpha \times \beta_j^{EA}$: indirect mediated effect that captures the genetic association of EA ($\beta_j^{EA}$) scaled by the correlation between INC factor and EA ($\alpha$)
- $\delta_j$: direct effect representing the genetic association of INC factor unexplained by EA (NonEA-INC).

In this model, the SNPs that are associated with EA but not associated with income (corresponding to the blue part in **Extended Data Fig. 2a**) are SNPs whose direct effects on income ($\delta_j$) are strong enough to offset their indirect effects ($\alpha \times \beta_j^{EA}$). These are SNPs with effects whose signs are discordant across EA and NonEA-INC. On the other hand, SNPs with null association for NonEA-INC imply that there will be a perfect genetic correlation between income and EA within such SNPs (corresponding to the orange part of **Extended Data Fig. 2a**). It is important to note that this model does not imply any causal direction between the variables. The above decomposition always holds for any given GWAS effects from two traits.

We estimated this model using genomic SEM[11], which essentially involves estimating the non-EA genetic association of INC factor ($\delta_j$) and the correlation between EA to income ($\alpha$).

Instead of the default European reference panel from phase 3 of the 1000 Genomes Project[2] provided by genomic SEM, we used the HRC European reference panel to increase the SNP coverage. Genomic SEM uses a reference panel to align SNPs and obtain MAF estimates, which in turn are used to compute the per-allele effect sizes standardized with respect to the phenotype. As a result, 7,274,585 SNPs were included in the final output.

#### 4.3. Concordant and discordant sets

We classified the SNPs as concordant or discordant on the basis of the sign concordance of $\hat{\delta}_j$ and $\hat{\beta}_j^{EA}$ estimates. Out of 7,274,585 SNPs, 4,056.295 SNPs were classified as discordant (corresponding to 55.8%). The sign discordance here implies that the size of $\hat{\beta}_j^{INC}$ is much smaller than that of $\hat{\beta}_j^{EA}$.

To validate this result, we tested in an independent sample whether the SNPs with the sign discordance between EA and NonEA-INC had weaker genetic associations of income. We grouped the SNPs into two sets according to the sign concordance. We then estimated the partitioned heritability of income for these two groups of SNPs in unrelated individuals from the sibling subsample of the UKB.

More specifically, we used GCTA[20] to construct two genomic relatedness matrices (GRM), one with 681,049 discordant SNPs and the other with 537,607 concordant SNPs, all of which were from the HapMap3 set. Here the SNPs were stratified from the GWAS results excluding the UKB sibling sample and their close relatives. After removing the individuals whose occupational or household income was not available, we identified unrelated individuals from the sibling sample by applying *--grm-singleton 0.05*, using a merged GRM from the two GRMs. Then, the heritability was estimated each for occupational and household income, specifying the two GRMs separately in the model. The sample size was 12,689 and 16,972 for occupational and household income, respectively. As covariates, we included age, age$^2$, age$^3$, sex, dummies for survey year, and interactions between sex and the rest.

We confirmed that the heritability of income was contributed disproportionately more by the concordant SNPs and far less by the discordant SNPs. The concordant SNPs accounted for 70.0% of the heritability (*s.e.* = 19.5%) for occupational income and 75.3% for household income (*s.e.* = 15.8%). In both cases, only the heritability component for the concordant SNPs was statistically significant.

### 5. Polygenic score analyses of income

#### 5.1. Baseline polygenic prediction

We conducted a validation analysis based on polygenic prediction with 1KG-EUR-like individuals in the Swedish Twin Registry (STR), which was not included in our meta-analysis. We chose the STR as the main prediction cohort for its accurate income data collected from administrative data sources, which include individual, occupational, and household income (see the subsection below for the detail). In addition, we also used the UKB siblings (UKB-sib) and the Health and Retirement Study (HRS) from the US as prediction cohorts. For the UKB-sib, occupational and household income measures were available, while a self-reported individual income measure was available for the HRS.

We constructed polygenic indexes (PGI), using the meta-analysis results of income excluding a prediction cohort at a time, as well as a PGI based on the EA GWAS summary statistics in the same way for comparison. PGIs were created only with HapMap3 SNPs[12] as these SNPs are known to have good imputation quality and provide good coverage in 1KG-EUR-like samples. Furthermore, the SNPs were limited to those available in both income and EA summary statistics for the sake of precise comparison. We used the reference panel from the HRC. The details of QC for this panel can be found in the Supplementary Information of ref[17].

We derived PGIs based on a Bayesian approach implemented in the software LDpred2[21]. LDpred2 is an extension of LDpred[22], which adjusts for LD and computes individual SNP weights by using posterior means of LD-independent effect-size distributions. LDpred2 improves LDpred approach by 1) using a LD window based on genetic distances, which can better accommodate long LD regions and 2) allowing for Bayesian updating of $p$ (the proportion of causal SNPs) and $h^2$ (SNP heritability) parameters (called LDpred2-auto). As priors, we set 0.2 for $p$ and LDSC $h^2$ estimates for $h^2$ parameters. While the authors of LDpred2 recommend running LDpred2 genome-wide, we ran Ldpred2 per chromosome for its computational efficiency given that prediction results are barely different for a well-powered GWAS.

Since the STR sample was genotyped with three different platforms, which gave too few common HapMap3 SNPs after quality-control filters, we applied LDpred2 for the SNPs available in each batch and created PGIs for each batch. We then included indicators for these different batches in the prediction analyses.

In order to create PGIs for the UKB siblings, we re-conducted the GWAS of income and EA excluding the sibling sample as well as their close relatives (up to the third degree of relatedness). We then performed the meta analyses again.

For both the STR and the UKB siblings, we randomly chose one sibling from each family to avoid complications due to having relatives in the sample. We measured the prediction accuracy on the basis of the incremental $R^2$, which is the difference between the $R^2$ from a regression of the phenotype on the PGI and the baseline covariates and the $R^2$ from a regression on the baseline covariates only. We constructed confidence intervals for the incremental $R^2$ by bootstrapping the sample 1,000 times.

Because income typically contains substantial demographic variation, we pre-residualized the log of income for demographic covariates. Then, as baseline covariates, we only included top 20 genetic PCs and genotype batch indicators. Because income data was available for multiple years for the STR and the HRS, we residualized the log of income for age, $age^2$, $age^3$, sex, and interactions between sex and the age terms within each year and obtained the mean of residuals for each individual. For the UKB-sib, which only had cross-sectional data, we residualized the log of income for age, $age^2$, $age^3$, sex, dummies for survey year, and interactions between sex and the rest. For EA measure (years of education), we applied the same procedure for consistency while using dummies of birth year in place of the age terms.

##### 5.1.1. The STR income data description

*Individual income*
We used work income (including work-related benefits) for each year from 1990 to 2018 from Sweden's registry data. For the years of 1970, 1975 and 1985, the census data was also used. The sample was limited to individuals of at least 30 years of age at each point. A few extreme outliers were removed by trimming at the 99th percentile.

*Household income*
Household income data only existed for a few years in the late census data. Therefore, all the observations were taken from the 1990s. The same sample filtering and the outlier removal was applied.

*Occupational income*
Work income averages (including work-related benefits) per category of three-digit ISCO codes (*SSYK*, the Swedish version) were obtained per year from 2001 to 2016 from the population level data (except for 2014 where the occupation codes were missing). These were then matched to the corresponding year-code observations in the STR data.

#### 5.2. Within-family polygenic prediction

Genetic associations of SES traits are known to particularly suffer confounds due to indirect genetic effects such as genetic nurture and population stratification[23–25]. Recent studies[17,26] have shown that, by controlling for parental PGIs, the direct genetic effect can be isolated from the overall population effect captured by the PGI. In the case of EA, the direct genetic effect was shown to account for 30.9% of the overall predictive power of the EA PGI.

We followed the same approach to estimate the share of the direct genetic effect in the overall population effect captured by INC factor PGI. We imputed missing parental genotypes from sibling and parent-offspring pairs, using the tool *snipar*[27]. To apply the imputation algorithm, we prepared the data as follows. We first identified individuals with "White British' ancestry" and first-degree relatives based on the kinship coefficients (first-degree if > 0.177) provided by the UKB. We then used KING software[27] to infer the sibling and parent–offspring relations by specifying "--related-degree 1".

As inputs for snipar, we only used high quality common SNPs from the HapMap3 set as well as directly genotyped SNPs (imputation INFO score > 0.99 and MAF>1%). We first phased these SNPs from the imputed genotype data using eagle v2.4.1[28] with 1000 Genomes Phase 3 reference panel. We then used these phased SNPs as inputs for snipar to impute missing parental genotypes. This procedure resulted in 1,244,153 SNPs in total whose missing rate was lower than 1%.

Following the same procedure in Section 5.1, we created a PGI for INC factor by applying LDpred2 to this set of SNPs for the UKB sibling sample. For each individual in this sample, we also created parental PGIs using the imputed parental genotypes. Each PGI was then normalized to have a variance of 1. The phenotypes (the log of occupational income or household income) were residualized for age, age$^2$, age$^3$, sex, dummies for survey year, and interactions between sex and the age and year terms, as well as top 20 genetic PCs and genotype batch indicators. Then, the residualized phenotypes were also normalized to have a variance of 1 separately in males and females. As a result, regression estimates of PGI represent (partial) correlations, and their squares indicate proportions of phenotypic variance explained.

For the prediction analysis, we estimated the following regression model:

$$y_{ij} = \delta PGI_{ij} + \alpha(PGI_{p(j)} + PGI_{m(j)}) + \varepsilon_{ij}$$

where $y_{ij}$ is the phenotype of individual $i$ in family $j$, $PGI_{ij}$ the PGI, $PGI_{p(j)}$ the paternal PGI, $PGI_{m(j)}$ the maternal PGI, and $\varepsilon_{ij}$ the error term. $\delta$ represents the direct genetic effect of the PGI and $\alpha$ reflects indirect genetic effects and the effects of other genetic and environmental factors including the confounding due to non-random mating. The derivations in ref[17,26] show that this regression gives an unbiased estimate of $\delta$, while the population effect, denoted $\psi$, is then equal to $\delta + (1 + r_{am})\alpha$, where $r_{am}$ is the correlation between $PGI_{p(j)}$ and $PGI_{m(j)}$.

We fitted the above regression by OLS and clustered the standard errors by family to account for within-family dependence. We estimated $r_{am}$ using the correlation between siblings' PGI, which is equal to $(1 + r_{am})/2$ (see the supplementary note section 8 of ref[26]). For INC factor PGI, this was estimated to be 0.107. Given these estimates, we computed estimates for $\psi$ as well as the ratio of direct genetic effect to the population effect, $\delta / \psi$ . We then derived standard errors for the estimates of $\psi$ and $\delta / \psi$ by using the delta method.

### 6. Genetic correlation analysis

We estimated genetic correlations of INC factor, EA, and NonEA-INC with a wide set of traits, including socioeconomic, behavioral, and physical and mental health traits. We used LDSC to estimate the genetic correlations ($r_g$), using the pre-computed LD scores from the authors. and computed the difference in $r_g$ between EA and NonEA-INC. We derived the standard errors for the difference in $r_g$ on the basis of jackknife estimates, using the same default approach of LDSC. Then, the false discovery rate correction was applied to the resulting p-values.

The phenotypes included in this genetic correlation analysis are as follows: Subjective well-being[14], Parental lifespan[29], Cognitive performance[7], General risk tolerance[9], Chronotype[30], Sleep duration[30], Age of smoking initiation[31], Smoking persistence[31], Cigarettes per day[31], Drinks per week[31], Alcohol dependence[32], Height[33], BMI[33], Waist-to-hip ratio[33],

Blood pressure (in-house GWAS conducted in the UKB sample), Type 2 diabetes[34], Triglycerides[35], ADHD[36], Bipolar disorder[37], Schizophrenia[38], Autism spectrum[39], Anorexia nervosa[40], Obsessive compulsive disorder[41], Major depressive disorder[42], Anxiety disorder[43], Neuroticism[44], Stress-related disorder[45], Cannabis use disorder[46], Cross disorder[47]

### 7. Biological annotation

#### 7.1. Gene mapping

We used FUMA v1.5.2[13] with the default parameters to find genes implicated in INC factor GWAS, which used four mapping approaches to link the identified SNPs to protein-coding genes. First, the genes were mapped to the SNPs on the basis of physical proximity with a 10 kb window. Second, the genes were mapped according to expression quantitative trait locus (eQTL). We used the eQTL data from GTEx v8[48] and BRAINEAC[49]. Third, the genes were mapped based on significant chromatic interactions with the builtin chromatic interaction data. Fourth, we considered the genes that were statistically significant with Bonferroni correction from MAGMA gene-based association tests, which convert the mean chi-square of member SNPs into a gene-level test statistic.

#### 7.2. Tissue-specific enrichment analysis

We performed tissue-specific enrichment analyses using two approaches: LDSC-SEG[50] and MAGMA gene-property analyses[51]. First, we applied LDSC-SEG to estimate tissue or cell-type specific enrichment, using the pre-computed LD scores by the authors according to the gene expression annotations from Franke lab[52] and GTEx v6 data[48]. Second, MAGMA gene-property analysis was used to examine relationships between gene-level associations and tissue-specific gene expression profiles. The gene expression data were taken from GTEx v8.

### 8. GREML heritability estimation

We estimated the heritability of different income measures in the STR and UKB-sib samples with GREML from GCTA[20]. Such estimates are useful for gauging the maximum predictive power that could be achieved by the PGI. For both data sets, only the HapMap3 SNPs were included in the GRM and unrelated individuals were identified by applying *--grm-singleton 0.05*. The phenotypes were residualized (and averaged per individual for the STR) prior to the analyses in the same way from the prediction analyses. The results were reported in the **Supplementary Table 13** along with the LDSC heritability estimates from the GWAS meta-analysis results.

### 9. Phenome-wide association study

We explored the clinical relevance of the INC factor PGI for common diseases in the sibling sample of the UKB. We conducted a phenome-wide association study, using the in-patient electronic health records for 115 diseases with sex-specific sample prevalence no lower than 1%. We derived case-control status according to the phecode scheme by mapping the UKB's

ICD-9/10 records to phecodes (https://phewascatalog.org/phecodes, version 1.2)[53,54]. These ICD-9/10 records were collected from hospitalization, cancer, and death registries (as of May 2021).

We fitted a linear regression of case-control status on the INC factor PGI while controlling for the parental PGIs to specifically capture the direct genetic effects of income PGI. As covariates, we also included year of birth, its square term, and their interactions with sex, genotype batch dummies, and 20 genetic PCs. The standard errors were clustered by family.

In total, 14 diseases from various categories were significantly associated with the direct genetic effect of the INC factor PGI at false discovery rate < 0.05 (**Extended Fig. 4** and **Supplementary Table 27**). The results suggest that having a higher INC factor PGI can lead to lower risk for cardiovascular diseases, digestive issues, Type 2 diabetes, obesity, depression, tobacco use disorder, and musculoskeletal issues.

#### 10. Cohort acknowledgements

##### 10.1. ALSPAC (Avon Longitudinal Study of Parents and Children)

We are extremely grateful to all the families who took part in this study, the midwives for their help in recruiting them, and the whole ALSPAC team, which includes interviewers, computer and laboratory technicians, clerical workers, research scientists, volunteers, managers, receptionists and nurses. The UK Medical Research Council and Wellcome (Grant ref: 217065/Z/19/Z) and the University of Bristol provide core support for ALSPAC. This publication is the work of the authors and they will serve as guarantors for the contents of this paper. GWAS data was generated by Sample Logistics and Genotyping Facilities at Wellcome Sanger Institute and LabCorp (Laboratory Corporation of America) using support from 23andMe.

Please note that the study website contains details of all the data that is available through a fully searchable data dictionary and variable search tool" and reference the following webpage: http://www.bristol.ac.uk/alspac/researchers/our-data/. Ethical approval for the study was obtained from the ALSPAC Ethics and Law Committee and the Local Research Ethics Committees. Consent for biological samples has been collected in accordance with the Human Tissue Act (2004). Informed consent for the use of data collected via questionnaires and clinics was obtained from participants following the recommendations of the ALSPAC Ethics and Law Committee at the time. At age 18, study children were sent 'fair processing' materials describing ALSPAC's intended use of their health and administrative records and were given clear means to consent or object via a written form. Data were not extracted for participants who objected, or who were not sent fair processing materials.

Study data were collected and managed using REDCap electronic data capture tools hosted at the University of Bristol. REDCap (Research Electronic Data Capture) is a secure, web-based software platform designed to support data capture for research studies. DOI: 10.1016/j.jbi.2008.08.010

Pregnant women resident in Avon, UK with expected dates of delivery between 1st April 1991 and 31st December 1992 were invited to take part in the study. 20,248 pregnancies have been identified as being eligible and the initial number of pregnancies enrolled was 14,541. Of the initial pregnancies, there was a total of 14,676 foetuses, resulting in 14,062 live births and 13,988 children who were alive at 1 year of age. The total sample size for analyses using any data collected after the age of seven is therefore 15,447 pregnancies. Of these 14,901 children were alive at 1 year of age. Of the original 14,541 initial pregnancies, 338 were from a woman who had already enrolled with a previous pregnancy, meaning 14,203 unique mothers were initially enrolled in the study. As a result of the additional phases of recruitment, a further 630 women who did not enrol originally have provided data since their child was 7 years of age. This provides a total of 14,833 unique women (G0 mothers) enrolled in ALSPAC as of September 2021. G0 partners were invited to complete questionnaires by the mothers at the start of the study and they were not formally enrolled at that time. 12,113 G0 partners have been in contact with the study by providing data and/or formally enrolling when this started in 2010. 3,807 G0 partners are currently enrolled.

##### 10.2. CoLaus (Cohorte Lausannoise)

The authors would like to thank all the people who participated in the recruitment of the participants, data collection and validation, particularly Nicole Bonvin, Yolande Barreau, Mathieu Firmann, François Bastardot, Julien Vaucher, Panagiotis Antiochos, Cédric Gubelmann, Marylène Bay and Benoît Delabays.

##### 10.3. Croatia - Korcula

This research was funded by the Medical Research Council UK, the Croatian National Centre of Research Excellence in Personalized Healthcare grant (number KK.01.1.1.01.0010), and the Centre of Competence in Molecular Diagnostics (KK.01.2.2.03.0006).

##### 10.4. EGCUT (Estonian Genome Center, University of Tartu)

The authors wish to acknowledge the participants of the Estonian Biobank for their contributions.
The activities of the EstBB are regulated by the Human Genes Research Act, which was adopted in 2000 specifically for the operations of EstBB. Individual level data analysis in EstBB was carried out under ethical approval by the Research Ethics Committee of the University of Tartu (Approval number 288/M-18), using data according to release application 6-7/GI/33516 from the Estonian Biobank.

The Estonian Genome Center analyses were partially carried out in the High Performance Computing Center, University of Tartu. The authors also acknowledge support for the development of the infrastructure of the Estonian Genome Centre from the Estonian Research

Infrastructures Roadmap project No. SP1GI16442T “Estonian Centre for Genomics II”. The work of the Estonian Genome Center, University of Tartu was funded by the European Union through Horizon 2020 research and innovation program under grants no. 810645 and 894987, through the European Regional Development Fund projects GENTRANSMED (2014-2020.4.01.15-0012), MOBEC008, MOBERA21 and Estonian Research Council Grants PRG791 and PRG1291. We also acknowledge the Estonian Biobank Research Team responsible for data collection, genotyping, quality control and imputation, including Andres Metspalu, Lili Milani, Reedik Mägi, Mari Nelis, Georgi Hudjashov.

##### 10.5. FTC (Finnish Twin Cohort)

Phenotype and genotype data collection in the twin cohort has been supported by the Wellcome Trust Sanger Institute, the Broad Institute, ENGAGE – European Network for Genetic and Genomic Epidemiology, FP7-HEALTH-F4-2007, grant agreement number 201413, and the Academy of Finland (grants 264146, 308248, 312073, 336823, and 352792 to JKaprio).

##### 10.6. HUNT (Trøndelag Health Study)

The Trøndelag Health Study (HUNT) is a collaboration between HUNT Research Centre (Faculty of Medicine and Health Sciences, NTNU, Norwegian University of Science and Technology), Trøndelag County Council, Central Norway Regional Health Authority, and the Norwegian Institute of Public Health. The genotyping in HUNT was financed by the National Institutes of Health; University of Michigan; the Research Council of Norway; the Liaison Committee for Education, Research and Innovation in Central Norway; and the Joint Research Committee between St Olavs hospital and the Faculty of Medicine and Health Sciences, NTNU. Laxmi Bhatta and Ben M. Brumpton received support from the HUNT Center for Molecular and Clinical Epidemiology; Faculty of Medicine and Health Sciences, NTNU; The Liaison Committee for Education, Research and Innovation in Central Norway; and the Joint Research Committee between St Olavs Hospital and the Faculty of Medicine and Health Sciences, NTNU.

##### 10.7. iPSYCH

The iPSYCH consortium is supported by the Lundbeck foundation (grant nos. R1-2=A9118 and R155-2014-1724). The funders had no role in study design, data collection and analysis, decision to publish or preparation of the manuscript.

##### 10.8. LifeLines

The Lifelines Biobank initiative has been made possible by funding from the Dutch Ministry of Health, Welfare and Sport, the Dutch Ministry of Economic Affairs, the University Medical Center Groningen (UMCG the Netherlands), University of Groningen and the Northern Provinces of the Netherlands. The generation and management of GWAS genotype data for the Lifelines Cohort Study is supported by the UMCG Genetics Lifelines Initiative (UGLI). UGLI is partly supported by a Spinoza Grant from NWO, awarded to Cisca Wijmenga. The authors wish to acknowledge the services of the Lifelines Cohort Study, the contributing research centers delivering data to Lifelines, and all the study participants.

The following individuals contributed to the Lifelines study:
Raul Aguirre-Gamboa (1), Patrick Deelen (1), Lude Franke (1), Jan A Kuivenhoven (2), Esteban A Lopera Maya (1), Ilja M Nolte (3), Serena Sanna (1), Harold Snieder (3), Morris A Swertz (1), Peter M. Visscher (3,4), Judith M Vonk (3), Cisca Wijmenga (1)
(1) Department of Genetics, University of Groningen, University Medical Center Groningen, The Netherlands
(2) Department of Pediatrics, University of Groningen, University Medical Center Groningen, The Netherlands
(3) Department of Epidemiology, University of Groningen, University Medical Center Groningen, The Netherlands
(4) Institute for Molecular Bioscience, The University of Queensland, Brisbane, Queensland, Australia.

#### 10.9. MOBA (Norwegian Mother, Father and Child Cohort Study)

The Norwegian Mother, Father and Child Cohort Study (MOBA) is supported by the Norwegian Ministry of Health and Care Services and the Ministry of Education and Research. We are grateful to all the participating families in Norway who take part in this on-going cohort study.

We thank the Norwegian Institute of Public Health (NIPH) for generating high-quality genomic data. This research is part of the HARVEST collaboration, supported by the Research Council of Norway (#229624). We also thank the NORMENT Centre for providing genotype data, funded by the Research Council of Norway (#223273), South East Norway Health Authorities and Stiftelsen Kristian Gerhard Jebsen. We further thank the Center for Diabetes Research, the University of Bergen for providing genotype data and performing quality control and imputation of the data funded by the ERC AdG project SELECTionPREDISPOSED, Stiftelsen Kristian Gerhard Jebsen, Trond Mohn Foundation, the Research Council of Norway, the Novo Nordisk Foundation, the University of Bergen, and the Western Norway Health Authorities.

This work was supported by in part by Research Council of Norway through its Centre of Excellence funding scheme, grant number 262700.

#### 10.10. NEO (The Netherlands Epidemiology of Obesity Study)

The NEO study is supported by the participating Departments, the Division, and the Board of Directors of the Leiden University Medical Centre, and by the Leiden University, Research Profile Area 'Vascular and Regenerative Medicine'. The authors of the NEO study thank all participants, all participating general practitioners for inviting eligible participants, all research nurses for data collection and the NEO study group: Pat van Beelen, Petra Noordijk and Ingeborg de Jonge for coordination, laboratory and data management.

##### 10.11. NTR (Netherlands Twin Registry)

We warmly thank all twin and family members for their participation.

##### 10.12. QIMR (Queensland Institute of Medical Research)

Data collection funded from various grants from Australian NHMRC and US NIH.

##### 10.13. RS (Rotterdam Study)

The generation and management of GWAS genotype data for the Rotterdam Study (RS I, RS II, RS III) was executed by the Human Genotyping Facility of the Genetic Laboratory of the Department of Internal Medicine, Erasmus MC, Rotterdam, The Netherlands. The GWAS datasets are supported by the Netherlands Organisation of Scientific Research NWO Investments (nr. 175.010.2005.011, 911-03-012), the Genetic Laboratory of the Department of Internal Medicine, Erasmus MC, the Research Institute for Diseases in the Elderly (014-93-015; RIDE2), the Netherlands Genomics Initiative (NGI)/Netherlands Organisation for Scientific Research (NWO) Netherlands Consortium for Healthy Aging (NCHA), project nr. 050-060-810. We thank Pascal Arp, Mila Jhamai, Marijn Verkerk, Lizbeth Herrera and Marjolein Peters, MSc, and Carolina Medina-Gomez, MSc, for their help in creating the GWAS database, and Karol Estrada, PhD, Yurii Aulchenko, PhD, and Carolina Medina-Gomez, MSc, for the creation and analysis of imputed data.

The Rotterdam Study is funded by Erasmus Medical Center and Erasmus University, Rotterdam, Netherlands Organization for the Health Research and Development (ZonMw), the Research Institute for Diseases in the Elderly (RIDE), the Ministry of Education, Culture and Science, the Ministry for Health, Welfare and Sports, the European Commission (DG XII), and the Municipality of Rotterdam. The authors are grateful to the study participants, the staff from the Rotterdam Study and the participating general practitioners and pharmacists.

##### 10.14. SHIP (Study of Health in Pomerania)

SHIP is part of the Community Medicine Research net of the University of Greifswald, Germany, which is funded by the Federal Ministry of Education and Research (grants no. 01ZZ9603, 01ZZ0103, and 01ZZ0403), the Ministry of Cultural Affairs as well as the Social Ministry of the Federal State of Mecklenburg-West Pomerania, and the network 'Greifswald Approach to Individualized Medicine (GANI_MED)' funded by the Federal Ministry of Education and Research (grant 03IS2061A). Genome-wide data have been supported by the Federal Ministry of Education and Research (grant no. 03ZIK012) and a joint grant from Siemens Healthineers, Erlangen, Germany and the Federal State of Mecklenburg- West Pomerania. The University of Greifswald is a member of the Caché Campus program of the InterSystems GmbH.

##### 10.15. STR (Swedish Twin Registry)

The Swedish Twin Registry is managed by Karolinska Institutet and receives funding through the Swedish Research Council under the grant no 2017-00641. Genotyping was performed by the SNP&SEQ Technology Platform in Uppsala (www.genotyping.se). The facility is part of the National Genomics Infrastructure supported by the Swedish Research Council for Infrastructures and Science for Life Laboratory, Sweden. The SNP&SEQ Technology Platform is also supported by the Knut and Alice Wallenberg Foundation.

##### 10.16. UKHLS (Understanding Society)

University of Essex, Institute for Social and Economic Research. (2022). Understanding Society: Waves 1-12, 2009-2021 and Harmonised BHPS: Waves 1-18, 1991-2009. [data collection]. 17th Edition. UK Data Service. SN: 6614, http://doi.org/10.5255/UKDA-SN-6614-18.
