## Supplementary material for "Associations between common genetic variants and income provide insights about the socioeconomic health gradient": Frequently Asked Questions

### Frequently Asked Questions (FAQs)

This document provides information about the study:

#### Table of contents:

**1. Summary** **3**
**2. The current study** **5**
2.1. What is the purpose of this study? 5
2.2. What did you do in this paper? 6
2.3. How was income measured in the current study? 7
2.4. Did you find the gene(s) for income? 7
2.5. Are the genetic variants associated with income in your study also associated with other outcomes? 9
2.6. How good is your polygenic index? 9
2.7. What does your study not mean? 10
2.8. Is this the first GWAS on income? 11
**3. Implications of the study** **11**
3.1. Does this study show that an individual's level of income is determined, or fixed, at conception? 11
3.2. Can your polygenic index be used to predict how well someone will do in life? 12
3.3. Can your polygenic index be used for research studies in non-European-ancestry populations? 13
3.4. What policy lessons do you draw from this study? How could society benefit from this work? 15
3.5. Could this kind of research lead to discrimination against, or stigmatization of, people with the relevant genetic variants? If so, why conduct this research? 16
**4. Background** **18**
4.1. What does it mean to say that income is "heritable"? 18
4.2. What is a GWAS? Are the genetic variants identified in a GWAS "causal"? 18
4.3. What is a polygenic index? 21
4.4. Where can I learn more about social science genetics? 22
**5. Appendices** **24**
5.1. Quality control measures 24
5.2. Additional reading and references 26

### 1. Summary


- We investigated associations between genetics and income and explored how these are linked to health outcomes. By analyzing data from 668,288 individuals in 12 affluent countries who carried genotypes most similar to the EUR reference panel of the 1000 Genomes dataset (1KG-EUR), we discovered that different measures of income share common genetic associations. These associations do not imply a direct genetic determination of income, but probably reflect the ways in which a particular society may reward certain genetic predispositions. We identified 162 genetic regions associated with these income measures, 88 of which were previously unknown to be linked to income. The effect of each individual genetic region is very small.
- We observed substantial heterogeneity in the genetic architecture of income across cohorts and non-perfect genetic correlations of income across sexes. This underlines that the genetic associations we report here are averages across different groups and environments that should not be interpreted as fixed or universal.
- By creating a polygenic index based on our genetic association results, we were able to capture 1 - 4% of the variation in income measures among 1KG-EUR-like individuals. The predictive power of the polygenic index decreased by ~75% when accounting for the polygenetic indexes of an individual's parents, indicating that assortative mating magnifies the predictive power of the polygenic index relative to the causal effects of the genetic variants and/or that environmental factors correlated with the family account for a substantial part of the link between genetic variants and income.
- Our analysis revealed that associations between higher income and better health outcomes are partly due to common genetic factors linked to both. Genetic effects associated with higher income correlate with lower BMI, blood pressure, type-2 diabetes, depression, and reduced stress-related disorders. Interestingly, genetic components of income not shared with educational attainment are related to better mental health, but reduced physical health benefits and increased risky behaviours such as drinking and smoking.
- We found genetic correlations between a range of psychiatric disorders and both the overall genetic effects associated with income and those specific to income - those not overlapping with educational attainment. Notably, for certain disorders - schizophrenia, autism, and obsessive-compulsive disorder - these correlations were positive when considering all genetic effects associated with income and educational attainment, and negative for those unique to income. This may indicate that the educational system may better accommodate individuals with these disorders than the labor market does, or that

talents associated with these genetic risks are advantageous in school but not in the labor market.

- It is of paramount importance to stress that our study does not provide any basis for asserting inherent superiority or accepting social inequality as an inevitability grounded in genetics. We firmly reject such notions on scientific and ethical grounds. Instead, our findings demonstrate that genetic endowments play a role in inequality and that their effects are influenced by environmental and societal factors that are susceptible to change. Moreover, our study results should not be used for making comparisons between different groups or predicting individual-level outcomes. Discrimination cannot be justified based on genetic associations.
- In summary, our study sheds light on the complex interplay between genetics, socio-economic status, and health outcomes. It underscores the importance of environmental and societal factors in these relationships, and supports the imperative for fairness in access to opportunities and resources for all individuals.

### 2. The current study

#### 2.1. What is the purpose of this study?

Understanding the causes of social mobility and in particular the structural sources of inequality is of fundamental importance both as a matter of science and of social policy.[2] Poverty and economic deprivation are major risk factors for mental and physical diseases,[3] lower life expectancy,[4] and lower well-being.[5] Furthermore, socio-economic status (SES) is important to health not only for those in poverty, but at all levels of SES.[6] It has long been recognized that parental SES is a major determinant of a child's expected trajectory in terms of cognitive and non-cognitive skill development, behaviors,[7] educational attainment,[8] career prospects, and adult income.[9] In other words, differences in SES are partially transmitted across generations. At the same time, education, income, personality, cognitive abilities, and occupational choices are all heritable to some extent and parents pass on both their environments and their genes to their offspring.[10–13] Consequently, disentangling the effects of behavior, environment, and genetics on income poses a substantial scientific challenge, yet remains important for comprehending social mobility and gaining insights into the intricate relationships between income and health.

Yet, the scientific possibilities to do so are limited. In the past, the primary tools to disentangle the effects of a parent's genes from parental environment were adoption studies[14] and children-of-twin studies.[15] However, few samples of this type exist, those that do are typically small, and these naturally occurring experiments are rarely representative of the entire range of environments. Furthermore, these datasets do not allow the investigation of any interactions between environments and specific biological pathways. As a result, scientific insights are still very limited about why and how social inequalities tend to persist within families throughout generations, why and how these inequalities translate into differences in health and mortality, and what the most effective ways are to help disadvantaged individuals.

With the advent of well-powered genome-wide association studies (GWAS) on socio-economic indicators such as educational attainment and income, new opportunities have emerged to help address these challenges. Polygenic indices derived from GWAS offer new approaches to investigate the contributions of direct genetic effects and environmentally mediated mechanisms in intergenerational social mobility, for example in samples of trios comprising mother, father, and child.[16] Additionally, well-powered GWAS summary statistics for a wide range of traits enable the exploration of genetic effects shared between income and health outcomes, with the potential to unveil previously unknown relationships. They also facilitate studies investigating interaction effects between genetic and environmental factors. Furthermore, incorporating polygenic indices from robust GWAS on income can help control for potential genetic confounds and enhance the statistical power of social scientific studies on income and social mobility.[17,18]

The goal of our study was to provide additional insights into common genetic variants associated with income, to shed light on potential societal biases towards certain genetic predispositions, and to gain information about the intricate relationships between health and socio-economic status. In particular, we used advanced statistical methods to disentangle the genetic associations with income and educational attainment (EA) and leveraged them to gain insights into distinct associations of these two major indicators of SES with health outcomes. We are sharing the GWAS summary statistics of our study with the broader research community. With this new data, researchers will be able to study a variety of important questions that will be informative about the structural sources of inequality and their relationships with health.

#### 2.2. What did you do in this paper?

We conducted a GWAS (What is a GWAS?) of income in a sample of over 600,000 participants. To construct such a large sample, we collected 32 datasets from 12 countries. All of these datasets have surveyed and genotyped their research participants. We considered four income measures (individual, household, occupational, parental). We conducted a GWAS for each income measure and combined the results.

We then used the findings of our GWAS to perform additional analyses that explored:

1. the environmental heterogeneity in the genetic factor for income, by comparing the results by sex and by countries,
2. similarities and differences of our results with the most recent GWAS for educational attainment,
3. the predictive power of the polygenic index for income as well as the influence of family environment, and

4. the relation with other behavioral and health-related phenotypes.

#### 2.3. How was income measured in the current study?

Our GWAS used four measures of income: individual, occupational, household, and parental income.

Individual income is the most direct measure of the consumption and savings opportunities that a person has. Individual income is the result of various factors including achieved qualifications (e.g., education, learned occupation, experience), personal characteristics (e.g., leadership, cognitive skills, consciousness), the demand and supply for these qualifications and characteristics in the labor market, and personal choices about labor supply (e.g., due to personal preferences, and decisions about division of labor among household members).

However, most large datasets that contain genetic information do not have measures of individual income. To address this challenge, we used three additional measures of income (household income, parental income, occupational income) that are all genetically highly correlated (**Fig 1a**). Household income was measured by questionnaires. Parental income (only available in the Danish iPsych sample) was derived from national registries, calculating the average income of mothers and fathers between ages 30 and 55. Occupational income was derived from standardized occupational codes of participants. Using the national statistics based on the same occupational coding, we imputed the average income within occupational codes for each participant.

#### 2.4. Did you find the gene(s) for income?

No.

We did not find "the gene for" income. We identified many genetic variants that are associated with income. Although it was once believed that scientists would discover numerous one-to-one associations between genes and outcomes, we have known for many years that the vast majority of human traits and other outcomes are complex and are influenced by many (thousands or even tens of thousands of) genes, each of which alone tends to have a small influence on the relevant outcome.[20]

Although we did find several genes that are associated with income, we believe that characterizing these as "genes for income" is likely to mislead, for many reasons.

First, a large part of the variation in people's income is accounted for by social and other environmental factors, not by additive genetic effects. "Genes for income" might be read to imply, incorrectly, that genes are the strongest predictor of variation in income — this is not the case.

Second, the genetic variants that are associated with income are also associated with many other things (only some of which we identify in this study, see, for example, our results on the links between income and health). These variants are no more "for" income than for the other outcomes with which they are associated (e.g., educational attainment, BMI, Alzheimer's disease, HDL cholesterol levels).

Third, each individual genetic variant captures only a tiny part of the variance in income (less than 0.011%). Our results suggest that hundreds, or even thousands, of genetic variants are associated with income, but each of them considered by itself has only a tiny effect. The phrase "genes for income" might misleadingly imply large effects of specific genes, but these effects do not exist.

Fourth, environmental factors can increase or decrease the impact of specific genetic variants. Put differently, even if a genetic variant is associated with higher or lower levels of income *on average*, it may have a much larger or smaller effect depending on social and environmental conditions. We illustrate this by calculating the *average* genetic correlation of income across the various datasets included in our meta-analysis. For individual income, this average genetic correlation across samples is only 0.45, suggesting a high degree of heterogeneity in the genetic architecture of income across different environments.

Finally, genes do not affect income directly. Rather, their influence works via social and environmental channels that are subject to change. For example, our results suggest that the associations between genes and income work partly through educational attainment which can be influenced by policy interventions. Furthermore, the predictive power of the polygenic index decreased by ~75% when accounting for the genetic indexes of an individual's parents, indicating that family environments account for a substantial part of the link between genetic variants and income.

##### 2.5. Are the genetic variants associated with income in your study also associated with other outcomes?

Yes.

##### 2.6. How good is your polygenic index?

Our polygenic index captures approximately 1 - 4% of the variance in different income measures in three hold-out samples from the United Kingdom, the United States, and Sweden (see also the FAQ section What is a polygenic index?). This is good enough for important research purposes such as exploring the relationships between health and income but is useless for (misguided) attempts to "predict" individual-level outcomes from genetic data (see section 3.2 below).

Note that our estimates of the "SNP heritability" of income (Supplementary Table 13) suggest a cohort-specific upper bound of 10~13% for the potential accuracy of polygenic indices for income in the future, as GWAS sample becomes larger (see the FAQ What is a GWAS?). Even a polygenic index of ~10% would not be accurate enough to make meaningful statistical predictions at the individual level — see Figure 3 in [18]. Furthermore, our prediction analyses suggest that approximately three-fourths of the signal our polygenic index picks up is actually due to environmental effects that are correlated with genes, such as the rearing environment provided by one's parents. Even the remaining effects are partially mediated by environmental channels such as educational attainment. This finding clearly illustrates what we said here earlier — polygenic indices are not a clean way to separate biological and environmental influences. **Genetic effects on socioeconomic outcomes such as income do not exist in a vacuum — they are shaped by environmental conditions that keep evolving and that are not equal for everyone.**

##### 2.7. What does your study *not* mean?

Genetic research has a long history of being misinterpreted and misused to argue that social inequality is inevitable, that social programs designed to improve people's lives are bound to fail and that some people are "naturally" inferior to other people.[22] **We wholeheartedly reject these claims on both scientific and moral grounds.**

A high or low polygenic index should not be interpreted to mean that someone is destined or determined to show a particular characteristic. It is not a "fortune teller", nor a pure measure of someone's genetic "endowment." It is just one of many factors that matter. By way of analogy, having high cholesterol makes it more likely that an individual will have a heart attack, but it does not determine that outcome — lots of people have high cholesterol but don't have a heart attack, and you can take steps to prevent a heart attack if you are at high risk. Similarly, a high polygenic index means that an individual has a slightly higher probability of obtaining a higher income, but that higher probability does not mean destiny.

Genetic associations with income do not mean that the environment does not make a difference. In fact, polygenic indices such as ours partly capture what most people would consider as environmental effects. If, for example, parents with particular genetic variants are more likely to live in a wealthy neighborhood with good schools, and if good schools make it more likely that children will get good jobs later in their lives, then this means that their children's polygenic indices for income will be correlated with the quality of schools they attended. Our analyses show that roughly three-fourths of the signal our polygenic index for income picks up is due to such indirect genetic effects or other environmental influences that just happen to be correlated with genes.

Genetic associations with income do not mean that interventions or policy reforms designed to combat inequalities are bound to fail. For example, interventions such as the Perry Preschool Project have shown that access to high-quality education is an effective way to improve lifetime outcomes, including a higher likelihood of getting a good job that pays well.

**The existence of genetic associations with income within a group of people does not tell us anything about whether there are average differences between racial or ethnic**

**groups, or why such differences, if they are observed, occur.** This is an important point, because racist and classist ideas about the allegedly "inferior" character of people of color and the poor have been used to justify eugenic policies.[23–25] Nothing about this study gives any sort of empirical support to these ideas.

##### 2.8. Is this the first GWAS on income?

No.

Two previous GWAS have looked at household income in the UK Biobank. These studies uncovered many interesting findings that help to illuminate the complex relationships between socioeconomic conditions and health.[26,27] The current study is based on a substantially larger sample size that we obtained by pooling data from 32 different samples across 12 economically advanced countries and three continents. The larger sample size allows us to discover additional genetic variants associated with income, yields a better-performing polygenic index, and allows more precise estimates of the genetic correlations of income with other traits. We also used a multivariate statistical approach that helps us to parse genetic associations with income that are also related with educational attainment and those that are not. We used the results of the latter analyses to gain novel insights into partly divergent health-implications of educational attainment and income.

#### 3. Implications of the study

##### 3.1. Does this study show that an individual's level of income is determined, or fixed, at conception?

No.

**Social and other environmental factors are the main drivers of variation in income.** However, even if it were true that genetic factors accounted for *all* of the differences among individuals in income, it would *still* not follow that an individual's income is "determined" at conception. There are at least three reasons for this:

First, some (if not all) genetic effects may operate through environmental channels.[28] Our study clearly illustrates the relevance of educational attainment for income, and education can be changed through environmental interventions (e.g., policy).

Second, even if the genetic associations with income operated entirely through non-environmental mechanisms that are difficult to modify (such as direct influences on the formation of neurons in the brain and the biochemical interactions among them), powerful environmental interventions could still change these genetic relationships. In a famous example suggested by the economist Arthur Goldberger, even if all variation in unaided eyesight were due to genes, there would still be enormous benefits from introducing eyeglasses.[29] Similarly, policies that guarantee a minimum wage or a basic income have incontrovertible relevance for living standards and the overall distribution of income.

##### 3.2. Can your polygenic index be used to predict how well someone will do in life?

No (see also the FAQ sections What is a polygenic index? and How good are your polygenic indices?).

The figure below illustrates the statistical reason why the polygenic index cannot be used to predict how well someone will do in life. It shows data from the Health and Retirement Study, one of our replication samples in our paper. The *x*-axis plots the values of the polygenic index for income among 6,171 individuals. The values of the index in that sample were standardized to have a mean of zero and a standard deviation of one. The *y*-axis plots values of log self-reported income, controlling for demographic variables (see the figure note). Each dot in the figure represents the combination of the polygenic index and income for one specific person in the sample. Despite the small, positive relationship between the index and income in the sample (indicated by the dotted red regression line), it can be seen clearly that the values of the index are not very informative about the income of any specific individual in the sample. In fact, a very broad range of income values is observed even for people who have very low or very high values of the polygenic index (e.g., those that are two standard deviations away from the mean).

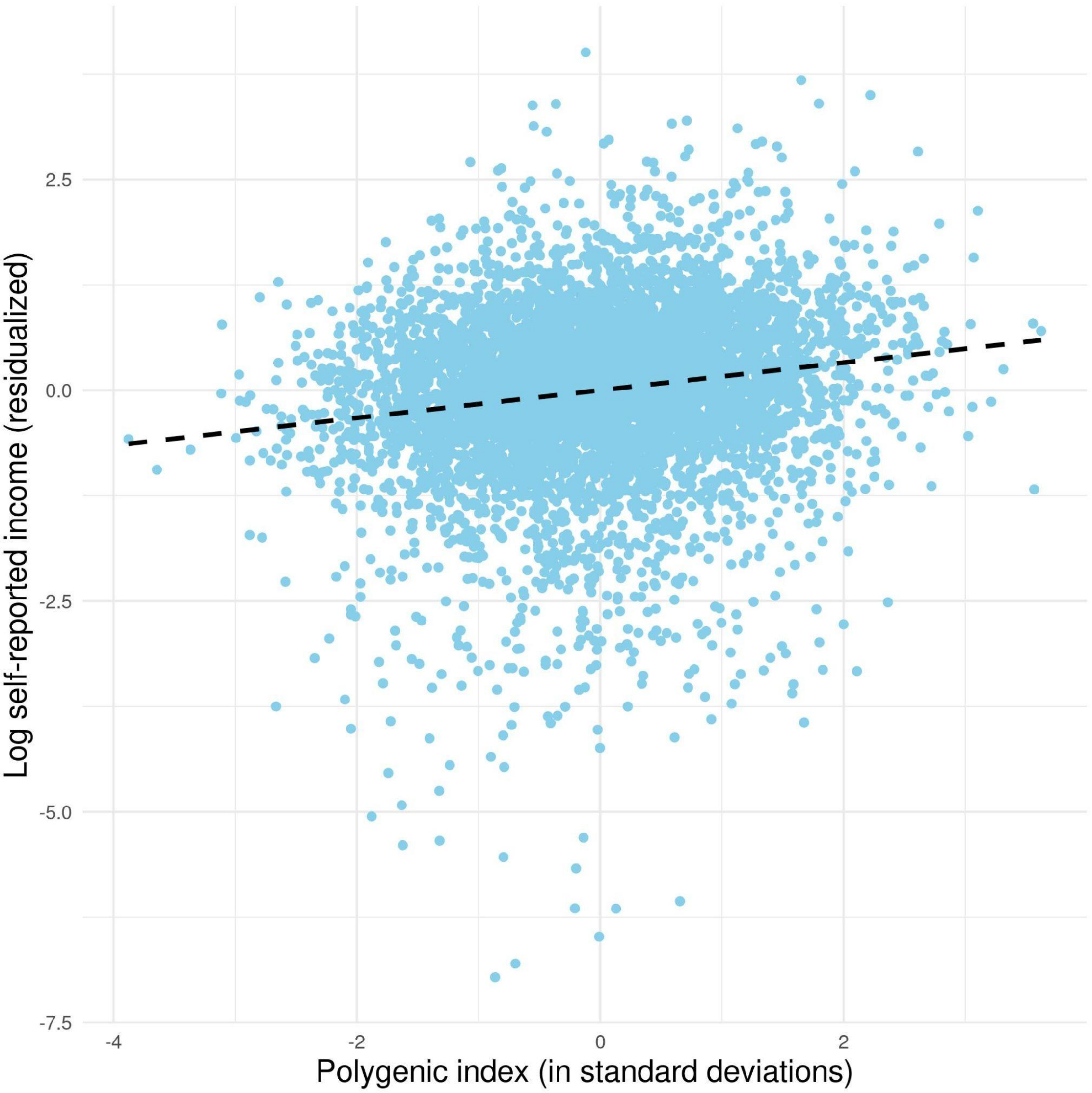


Note: The figure presents a scatter plot for the Health and Retirement Study (*N*=6,171). The x-axis shows standardized values of the polygenic index for log self-reported income, which was constructed as follows: within each wave, the log income was regressed on demographic variables (sex, age, $age^2$, $age^3$, and the interactions between sex and the age terms) and genetic control variables (top 20 genetic principal components and genotyping batches). Then, the mean of the residuals from the regressions was obtained for each individual, which was then standardized. The dotted line is a regression line with slope 0.172 ($p = 5.3\times\ 10^{-42}$).

#### 3.3. Can your polygenic index be used for research studies in non-European-ancestry populations?

Only in a very limited way.

As a practical matter, it is possible to calculate a polygenic index for any individual for whom genome-wide data are available, but the polygenic index is expected to be much less accurate for non-European-ancestry populations.[30] For example, a polygenic index for educational attainment constructed from a GWAS in over 3 million 1KG-EUR-liked individuals captures 12.0% of the variance in years of schooling among individuals with similar ancestries in the US Health and Retirement Study. The same index captures only 1.3% of the variance in years of schooling among African-Americans in the same sample.[31] For income, we expect a similarly strong attenuation of accuracy.

Our study was conducted only using 1KG-EUR-like samples of individuals (see the FAQ section Quality control measures). This choice was based primarily on data availability since the vast majority of all genotyped samples at the time of our study are 1KG-EUR-like (colloquially referred to as "European ancestry" below).[39] The set of SNPs that are associated with income in individuals of European ancestry is unlikely to overlap perfectly with the set of SNPs associated with income sampled in other parts of the world. Even if a given SNP is associated in both ancestry groups, the effect size—in other words, the strength of the association—will almost surely differ, partly because linkage disequilibrium (LD) patterns (i.e., the correlational structure of the genome) vary by ancestry.[32] Thus, some variants may be associated with income because the variant is in LD (i.e., correlated) with a variant elsewhere in the genome that causally affects income (What is a GWAS? Are the genetic variants identified in a GWAS "causal"?). If the strength of the correlation is greater in one ancestry group than in another, then the size of the association will be larger in that ancestry group. Moreover, even if LD patterns were similar in each ancestry group, the association may differ in different groups because the social and environmental conditions differ (What is a polygenic index?). Different socio-economic systems may assign differing values to certain genetic predispositions, impacting income potential. Thus, a genetic predisposition that increases income generation in one group or society may not have the same effect in another. The fact that there are differences across ancestry groups in the set of associated SNPs and their effect sizes has two important implications.

First, it means that the *polygenic indices of individuals from different ancestry groups cannot be meaningfully compared*. A recent paper[30] illustrated this point in the context of polygenic indices for predicting height; in the sample analyzed in that paper, polygenic indices for height for individuals of European ancestry are on average larger than those of South Asian ancestry which in turn are larger than those of African ancestry. In actuality, however, populations of African ancestries represented by the sample have similar heights to populations of European ancestries, and both African and European populations tend to be taller than South Asian populations.

Second, while polygenic indices may be used to study differences across individuals within a sample of people of non-European-ancestries, the accuracy of the index will be much smaller than that in a sample of people of European ancestries. Such an attenuation of predictive power has been repeatedly found in prior work.[33–36]

Third, the existence of genetic associations with income within a group of people (in our study, individuals of European ancestries) does not tell us anything about whether there are average differences between racial or ethnic groups, or why such differences, if they are observed, occur. This is an important point, because racist and classist ideas about the allegedly "inferior" character of people of color and the poor have been used to justify eugenic policies.[23–25] Nothing about this study gives any sort of empirical support to these ideas.

For a more extensive, excellent discussion of these and related issues, see Graham Coop's blog post "Polygenic scores and tea drinking": https://gcbias.org/2018/03/14/polygenic-scores-and-tea-drinking/.

##### 3.4. What policy lessons do you draw from this study? How could society benefit from this work?

We do not make direct recommendations for policy in this paper.

However, our results show that how well people do in life depends to some degree the "genetic lottery"[38] (i.e., which particular combination of your parents' genes you inherited). No one decides to participate in these lotteries or has any influence on its outcome. Our within-family analyses demonstrate that random genetic differences between siblings influence how they earn and how healthy they are (**Extended Fig 4**). Although the overall effect sizes of these random genetic difference on income and health are small, the existence of them limits the extent to which people can take personal credit for the good things that happen to them and may inspire some humility. It also puts limits on how much people can be blamed for things that did not turn out in their favor. Furthermore, our results show that luck and agency (e.g., the choices people make, and how hard they work) cannot be clearly separated from one another — our polygenic index is associated with education and income, two measures that many may intuitively file under effort rather than luck. Our hope is that the insight that (genetic and social) luck matters may contribute to a greater sense of solidarity in society, more empathy for those who are unlucky, and a greater willingness to share among those who have the means to give. This is our preferred interpretation of our empirical results, which is in line with arguments put forward in moral philosophy[39,40] and with empirical studies that found that people are less willing to tolerate inequality that is due to luck rather than effort.[41–46] However, this is not the only way to interpret our results and different policy implications may be derived from different philosophical perspectives.[47]

In general, the widespread availability of genetic data and the increasing insights into how genetic factors are associated with observed differences between people creates new challenges for policy making across various domains, including insurance markets, labor markets, personalized medicine, and reproductive technologies. In many of these areas, societies and policy makers will have to make difficult and complex trade-offs involving the protection of human rights, including respect for autonomy, dignity, privacy, the right to science, the right to know or not to know about genetic results, as well as the feasibility of certain forms of insurance or possible improvements in health and well-being.[48] One of the authors of the current study, Koellinger, recently wrote a policy report for the European Commission that reviews some of these challenges for policy making. This report can be accessed here.

We would like to state explicitly and emphatically that the genetic association results reported in the current study should not be used for any form of individual selection or discrimination (e.g., in labor or insurance markets or in reproductive technologies); see the FAQ sections What does your study not mean?, Can your polygenic index be used to predict how well someone will do in life?, and Can your polygenic index be used for research studies in non-European-ancestry populations? .

##### 3.5. Could this kind of research lead to discrimination against, or stigmatization of, people with the relevant genetic variants? If so, why conduct this research?

Unfortunately, like a great deal of research—including, for instance, research identifying genomic variants associated with increased cancer risk—the results can be misunderstood and misapplied. This includes misuses that would discriminate against people who carry specific genetic variants (e.g., in insurance or labor markets). Nevertheless, for a variety of reasons, we do not believe that the best response to the possibility that useful knowledge might be misused is to refrain from producing that knowledge. Furthermore, we do not claim that genetically-informed studies are better than other empirical approaches. However, we briefly discuss some of the broad potential benefits of this research. We then describe what we take to be our ethical obligation as researchers conducting this work.

First, a benefit of conducting social science genetics research in ever larger samples is that doing so allows us to correct the scientific record. An important theme in earlier work by the SSGAC has been to point out that most existing studies in social-science genetics that report genetic associations with behavioral traits have serious methodological limitations, fail to replicate and are likely to be false-positive findings.[49–52] This same point was made in an editorial in *Behavior Genetics* (the leading journal for the genetics of behavioral traits), which stated that "it now seems likely that many of the published [behavior genetics] findings of the last decade are wrong or misleading and have not contributed to real advances in knowledge".[53] One of the most important reasons why earlier work has generated unreliable results is that the sample sizes were far too small, given that the true effects of individual genetic variants on behavioral traits are small.

Second, social science genetics also has the potential to correct the *social* record and thereby to help *combat* discrimination and stigmatization. Our study clearly demonstrates that (i) the vast majority of income inequality is due to environmental factors, and (ii) the genetic influences we identified are not deterministic. Instead, they also depend on environmental factors and work via environmental channels that can be influenced (e.g., via educational attainment).

Third, social science genetics research has the potential to yield many other benefits, as briefly summarized in the FAQ section Where can I learn more about social science genetics? and further explained in [18]. Foregoing this research necessarily entails foregoing these and any other possible benefits, some of which will likely be the result of serendipity rather than being foreseeable.

In sum, we agree with the U.K. Nuffield Council on Bioethics, which concluded in a report [54], p.114) that "research in behavioural genetics has the potential to advance our understanding of human behaviour and that the research can therefore be justified," but that "researchers and those who report research have a duty to communicate findings in a responsible manner." In our view, responsible behavioral genetics research includes a sound methodology and analysis of data; a commitment to publish all results, including any negative results; and the transparent, complete reporting of the methodology and findings in publications, presentations, and communications with the media and the public, including particular vigilance regarding what the results do—and do not—show (hence, this FAQ document).

### 4. Background

#### 4.1. What does it mean to say that income is "heritable"?

"Heritable" is a confusing word that is often misinterpreted to mean that heritable characteristics are determined by biology. This is not true. As we describe in our article in depth, heritable characteristics — including income — *can* be influenced by the environment, *can* be changed by interventions or policy reforms, and are *not* "destined" to develop in a particular way based just on a person's genes. "Heritable" means that, within the sample of people being studied, those who are more genetically different from each other tend to show more different characteristics.[55] We return to this point — that heritable characteristics are not biologically determined — in the FAQ sections What is a GWAS? Are the genetic variants identified in a GWAS "causal"? and What does your study not mean?.

Previous studies have measured income and similar traits in identical and fraternal twins, and found that identical twins are more similar in their income skills than fraternal twins are.[56] This result tells us that income has a heritable component. Using different data and a different approach (i.e., by quantifying whether observed genetic differences between individuals correspond to differences in income), we come to the same conclusion in our study. Because we now know that virtually all aspects of behavior and personality that differ between people are at least partly heritable, this result is not surprising.[12,57] The heritability of income should not be taken to mean that income is especially "genetic" in some way.

#### 4.2. What is a GWAS? Are the genetic variants identified in a GWAS "causal"?

The short answers to these questions are as follows, with links to more details listed below:

- GWAS (genome-wide association studies) systematically scan the entire human genome for associations with a trait of interest, one SNP at a time.
- Correlations between SNPs are ignored.
- Not all potentially relevant genetic variants are included in a GWAS, partly because they may not have been measured precisely enough or measured at all.

- How frequently a particular genetic variant occurs in a population varies across places, which implies that genes and environments are correlated.
- GWAS try to control for potential correlations between genes and environments, but most GWAS (including ours) do so only imperfectly.
- Genes often influence outcomes only indirectly via environmental and behavioral pathways that can be changed.
- **For all of the reasons above, genetic variants identified in a GWAS are generally not directly causal**.

In a genome-wide association study (GWAS), scientists look at genetic variants measured across the entire human genome to see whether any of them are, *on average*, associated with higher or lower levels of some outcome. Similar to other studies our analyses focus on the most common genetic variants—so called single-nucleotide polymorphisms (SNPs). SNPs are sites in the genome where single DNA base pairs commonly differ across individuals. SNPs usually have two different possible base pairs, or alleles. Although there are tens of millions of sites where SNPs are located in the human genome, GWASs typically investigate only SNPs that can be measured (or imputed) with a high level of accuracy. Currently, such procedures usually yield millions of SNPs that together capture the most common genetic variations across people.

GWAS has been a successful research strategy for identifying genetic variants associated with many traits and diseases, including body height[58–60], Alzheimer's disease[61,62], and schizophrenia[63,64]. It has also recently been used to identify genetic variants associated with a variety of health-relevant social science outcomes, such as the number of children a person has[65], happiness[66,67], and educational attainment[31,66,68].

Statistical geneticists would call a genetic variant causal if a *ceteris paribus* change in that variant at conception would lead to a different outcome (e.g., higher income). Note that this definition of causality requires no minimum effect size (i.e., the change in income due to the change in the genetic variant may so small that it is barely measurable), nor does it require an understanding of the mechanism (i.e., the genetic effect may be indirect and mediated by environmental responses, self-selection into environments and so forth).

GWAS identifies genetic variants that are associated with the outcome, but an observed association with a specific variant does not imply that the variant *causes* the outcome, for a variety of reasons. First, genetic variants are often highly correlated with other, nearby variants on the same chromosome. As a result, when one or more variants in a region causally influence an outcome (in that particular environment), many noncausal variants in that region may also be identified as associated with the outcome. When GWAS results are analyzed, researchers often emphasize results for the genetic variant in a region that showed the strongest evidence of association. This variant does not need to be the causal variant. In fact, the causal genetic variant may not have even been measured directly. For example, GWAS that focus on common SNPs would not be able to identify rare or structural genetic variants (e.g., deletions or insertions of an entire genetic region) that are causal, but they may identify SNPs that are correlated with these unobserved variants.

Second, the frequencies of many genetic variants vary systematically across environments. If those environmental factors are not accounted for in the association analyses, some of the associations found may be spurious. To use a well-known example,[69] any genetic variants common in people of Asian ancestries will be associated statistically with chopstick use, but these variants would not *cause* chopstick use; rather, these genetic variants and the outcome of chopstick use are both distributed unevenly among people with different ancestries. This is the problem of "population stratification". GWAS researchers have a number of strategies for addressing the challenges posed by population stratification (see the FAQ section Quality control measures).

Even in studies such as ours that attempt to address and correct for heterogeneity in genetic ancestry, allele frequencies may nonetheless vary systematically with environmental factors. For example, a genetic variant that is associated with higher income in the parental generation may have downstream effects on children's educational outcomes (e.g., through living in a neighborhood with good schools). This same genetic variant is likely to be inherited by the children of these parents, creating a correlation between the presence of the genetic variant in a child's genome and the extent to which the child was reared in an environment with specific characteristics. A recent study of Icelandic families showed that the parental allele that is *not* passed on to the parent's offspring is still associated with the child's educational attainment, suggesting that GWAS results for educational attainment partly represent these intergenerational pathways.[16] Our sibling analyses yield results that are consistent with this conclusion.

Third, variants' effects on an outcome may be indirect, so a variant that may be "causal" in one environment may have a diminished effect or no effect at all in other environments. For example, the nicotinic acetylcholine receptor gene cluster on chromosome 15 is associated with lung cancer.[70–72] From this observation alone we cannot conclude that these genetic variants cause lung cancer through some direct biological mechanism. In fact, it is likely that these genetic variants increase lung cancer risk through their effects on smoking behavior. In a tobacco-free environment, it is plausible that many of these associations would be substantially weaker and perhaps disappear altogether. Thus, even *if* we have credible evidence that a specific association is not spurious, it is entirely possible that the genetic variant in question influences the outcome through channels that we, in common parlance, would label environmental (e.g., smoking). Nearly forty years ago, the sociologist Christopher Jencks criticized the widespread tendency to mistakenly treat environmental and genetic sources of variation as mutually exclusive, see also [57]. As the example of smoking illustrates, it is often overly simplistic to assume that "genetic explanations of behavior are likely to be exclusively physical explanations while environmental explanations are likely to be social".[28]

##### 4.3. What is a polygenic index?

The results of GWAS can be used to create a "polygenic index" (often also called a polygenic score), an index composed of many genetic variants from across the genome. More precisely, GWAS results are used to create a *formula* for how to construct a polygenic index. Using this formula, a polygenic index can then be constructed for any individual with genome-wide data. Indeed, some of the value of GWAS is that the polygenic index it produces can be used in subsequent studies conducted in other samples.

Because a polygenic index aggregates the information from many genetic variants, it is more strongly associated with variation among individuals for the GWAS outcome than with any single genetic variant. Often, the polygenic indices that are most strongly associated with an outcome are those created using *all* common genetic variants (typically more than a million) studied in GWAS. The larger the GWAS sample is, the more precise the polygenic index constructed from the GWAS results will be. If GWAS samples were infinitely large, the polygenic indices constructed from those samples would be expected to capture as much of the variance of a trait in an independent sample as implied by the SNP heritability of a trait. For example, in our study, we estimated the SNP heritability for income to be approximately 10%. That means that a polygenic index constructed from an infinitely large GWAS on income would be expected to capture approximately 10% of the variance in income in an independent sample that was not included in the GWAS.[73,74]

Note that this does not mean that 10% of anyone's income is biologically determined (see the FAQ sections What does your study not mean? and Does this study show that an individual's level of income, education, or health is determined, or fixed, at conception?). It is important to understand that polygenic indices are not a "clean" way to separate biological from non-biological factors that contribute to differences between people's outcomes.[18] GWAS results are not entirely immune to unobserved (e.g., environmental) confounds, such as parenting, or neighborhood characteristics, and genetic influences are often conditional on and/or mediated by environmental channels (What is a GWAS? Are the genetic variants identified in a GWAS "causal"? and [75]). For example, a society that systematically discriminates against people of color would induce a correlation between all genes for skin or hair pigmentation and income. However, a change in that society that would eliminate such discrimination could make these genetic associations disappear. Since polygenic indices merely aggregate the effects that were estimated in a GWAS, they partly reflect currently existing social realities. Furthermore, polygenic indices may exhibit different predictive accuracy even among members of the same ancestry group that vary from each other in terms of sex or socioeconomic status.[76]

##### 4.4. Where can I learn more about social science genetics?

Two of the coauthors of the current study, Harden and Koellinger, published a review on social science genetics in *Nature Human Behavior* [18] that provides a comprehensive answer. In short, we believe that the social sciences are incomplete without genetics because most differences between people in terms of their behavior, preferences, achievements, and life events are at least to some extent "heritable" (see the FAQ section What does it mean to say that income is "heritable"?). Thus, integrating molecular genetics into the social sciences presents an opportunity to deliver richer, more precise answers to old questions in psychology, sociology, economics and related fields. Furthermore, the new type of data and study designs that molecular genetics allow will grant social scientists opportunities to ask new questions and to pursue new answers that would not have previously been feasible. We see our current study as an example for both of these claims (What did you do in this paper?).

The review by Harden and Koellinger also discusses the challenges of this work more broadly and provides several substantive examples of new insights afforded by integrating molecular genetic data into the social sciences (e.g., with respect to the intergenerational transmission of human capital, social mobility over the lifespan, human demography, gene-environment interactions, and the intersection between “social/behavioral” genetics and “disease” genetics).

### 5. Appendices

#### 5.1. Quality control measures

There are many potential pitfalls that could lead to spurious results in genome-wide association studies (GWAS). We took many precautions to guard against these pitfalls.

One potential source of spurious results is incomplete "quality control (QC)" of the genetic data. To avoid this problem, we used state-of-the-art QC protocols from medical genetics research.[77] We supplemented these protocols by developing and applying additional, more stringent QC filters.

Another potential source of spurious results is a confound known as "population stratification." To give a well-known illustration, suppose we were conducting a GWAS on the use of chopsticks.[69] People of Asian ancestries are more likely to use chopsticks than people of European ancestries. If we combined samples of Chinese and European ancestries and performed a GWAS that ignores ancestry, then we would find genetic associations for these variants. However, those associations would simply reflect the fact that allele frequencies vary across ancestry groups.

In our study we were extremely careful to correct for population stratification as much as possible. At the outset, we restricted the study to individuals of European ancestries. As is standard in GWAS, we also controlled for "principal components" of the genetic data in the analysis; these principal components capture the small genetic differences across ancestry groups within European populations, so controlling for them largely removes the spurious associations arising solely from these small ancestry differences.

After taking these steps to minimize bias stemming from population stratification, we conducted follow up analysis with ~15,000 pairs of siblings (~30,000 individuals). These "within-family" analyses break the link between genes and the family environment, thanks to the natural experiment of meiosis. During meiosis, the two copies of each parental chromosome are randomly combined and then separated to create a set of two gametes (e.g., two eggs or two sperm) each of which contains only one new, resampled copy of each chromosome. This process creates an almost infinite number of different DNA sequences that each parent could theoretically pass on to their children. The resulting genetic differences between full siblings and dizygotic twins are therefore random and independent from environmental factors that vary between families. Therefore, comparisons among biological siblings yield estimates for our polygenic index that are immune to genetic nurture, uncontrolled population structure, and other sorts of environmental influences that cannot be traced back to direct genetic effects. In these analyses, we found that approximately 75% of the signal of our polygenic index is due to such environmental confounds, while the remaining 25% is due to genetic effects that originate in people we studied. We call these latter effects causal, even though the pathways from genes to outcomes are complex and often work via environmental channels (e.g. education, interactions with peers).

#### 5.2. Additional reading and references

Note: Whenever possible, we included links to freely available versions of these references.


1. Demange, P. A. *et al.* Investigating the genetic architecture of non-cognitive skills using GWAS-by-subtraction. *bioRxiv* 2020.01.14.905794 (2020) doi:10.1101/2020.01.14.905794.
2. Piketty, T. Social Mobility and Redistributive Politics. *Q. J. Econ.* **110**, 551–584 (1995).
3. Wilkinson, R. G. & Marmot, M. *Social Determinants of Health: The Solid Facts*. (World Health Organization, 2003).
4. Stringhini, S. *et al.* Socioeconomic status and the 25× 25 risk factors as determinants of premature mortality: a multicohort study and meta-analysis of 1\textperiodcentered 7 million men and women. *Lancet* **389**, 1229–1237 (2017).
5. Stevenson, B. & Wolfers, J. Subjective Well-Being and Income: Is There Any Evidence of Satiation? *Am. Econ. Rev.* **103**, 598–604 (2013).
6. Adler, N. E. *et al.* Socioeconomic status and health. The challenge of the gradient. *American Psychology* **49**, 15–24 (1994).
7. Heckman, J. J. & Mosso, S. The Economics of Human Development and Social Mobility. *Annu. Rev. Econom.* **6**, 689–733 (2014).
8. Haveman, R. & Smeeding, T. The role of higher education in social mobility. *Future Child.* **16**, 125–150 (2006).
9. Acemoglu, D. Technical Change, Inequality, and the Labor Market. *J. Econ. Lit.* **40**, 7–72 (2002).
10. Plomin, R., DeFries, J. C., Knopik, V. S. & Neiderhiser, J. M. *Behavioral Genetics*. (Worth Publishers, 2012).
11. Okbay, A. *et al.* Polygenic prediction of educational attainment within and between families from genome-wide association analyses in 3 million individuals. *Nat. Genet.* **54**, 437–449 (2022).
12. Polderman, T. J. C. *et al.* Meta-analysis of the heritability of human traits based on fifty years of twin studies. *Nat. Genet.* **47**, 702–709 (2015).
13. Krapohl, E. & Plomin, R. Genetic link between family socioeconomic status and children's educational achievement estimated from genome-wide SNPs. *Mol. Psychiatry* **21**, 437–443 (2016).
14. Horn, J. M., Loehlin, J. C. & Willerman, L. Intellectual resemblance among adoptive adoptive and biological relatives: the Texas adoption project. *Behav. Genet.* **9**, 177–201 (1979).
15. D'Onofrio, B. M. *et al.* The role of the children of twins design in elucidating causal relations between parent characteristics and child outcomes. *J. Child Psychol. Psychiatry* **44**, 1130–1144 (2003).
16. Kong, A. *et al.* The nature of nurture: Effects of parental genotypes. *Science* **359**, 424–428 (2018).
17. DiPrete, T. A., Burik, C. A. P. & Koellinger, P. D. Genetic instrumental variable regression: Explaining socioeconomic and health outcomes in nonexperimental data. *Proceedings of the National Academy of Sciences of the United States of America* **115**, E4970–E4979 (2018).
18. Harden, K. P. & Koellinger, P. D. Using genetics for social science. *Nature Human Behaviour* **4**, 567–576 (2020).
19. Visscher, P. M. *et al.* 10 Years of GWAS Discovery: Biology, Function, and Translation. *Am. J. Hum. Genet.* **101**, 5–22 (2017).

20. Chabris, C. F., Lee, J. J., Cesarini, D., Benjamin, D. J. & Laibson, D. I. The fourth law of behavior genetics. *Curr. Dir. Psychol. Sci.* **24**, 304–312 (2015).
21. Dougherty, C. Why are the returns to schooling higher for women than for men? *J. Hum. Resour.* **XL**, 969–988 (2005).
22. Kevles, D. J. *In the Name of Eugenics: Genetics and the Uses of Human Heredity*. (Harvard University Press, 1995).
23. Moore, T. O. Stamped from the Beginning: The Definitive History of Racist Ideas in America, by Ibram X. Kendi. *Black Scholar* **48**, 71–74 (2018).
24. Crouch, C. Molly Ladd-Taylor: Fixing the Poor: Eugenic Sterilization and Child Welfare in the Twentieth Century. Preprint at https://idp.springer.com/authorize/casa?redirect_uri=https://link.springer.com/article/10.1007/s10964-020-01218-w&casa_token=vQPCN-yroYoAAAAA:iuJEb2oPmHxKeorxPy59lLCGUtrAXqo1kmyB9LYUgXDhht4O3163bqZieggTcM1Ec_R-TVKRk5X8U2w (2020).
25. Carlson, E. A. Three Generations, No Imbeciles: Eugenics, the Supreme Court, and Buck v. Bell. By Paul A. Lombardo. Baltimore (Maryland): Johns Hopkins University Press. $29.95. xv 365 p.; ill.; index. 978‑0‑8018‑9010‑9. 2008. *The Quarterly Review of Biology* vol. 84 178–180 Preprint at https://doi.org/10.1086/603452 (2009).
26. Hill, W. D. *et al.* Molecular genetic contributions to social deprivation and household income in UK Biobank. *Curr. Biol.* **26**, 3083–3089 (2016).
27. Hill, W. D. *et al.* Genome-wide analysis identifies molecular systems and 149 genetic loci associated with income. *Nat. Commun.* **10**, 5741 (2019).
28. Jencks, C. Heredity, environment, and public policy reconsidered. *Am. Sociol. Rev.* **45**, 723–736 (1980).
29. Goldberger, A. S. Heritability. *Economica* **46**, 327–347 (1979).
30. Martin, A. R. *et al.* Clinical use of current polygenic risk scores may exacerbate health disparities. *Nat. Genet.* **51**, 584–591 (2019).
31. Lee, J. J. *et al.* Gene discovery and polygenic prediction from a genome-wide association study of educational attainment in 1.1 million individuals. *Nat. Genet.* **50**, 1112–1121 (2018).
32. 1000 Genomes Project Consortium *et al.* A global reference for human genetic variation. *Nature* **526**, 68–74 (2015).
33. Domingue, B. W., Belsky, D., Conley, D., Harris, K. M. & Boardman, J. D. Polygenic Influence on Educational Attainment: New evidence from The National Longitudinal Study of Adolescent to Adult Health. *AERA Open* **1**, 1–13 (2015).
34. Vassos, E. *et al.* An Examination of Polygenic Score Risk Prediction in Individuals With First-Episode Psychosis. *Biol. Psychiatry* **81**, 470–477 (2017).
35. Domingue, B. W. *et al.* Mortality selection in a genetic sample and implications for association studies. *International Journal of Epidemiology* vol. 46 1285–1294 Preprint at https://doi.org/10.1093/ije/dyx041 (2017).
36. Belsky, D. W. *et al.* Development and evaluation of a genetic risk score for obesity. *Biodemography Soc. Biol.* **59**, 85–100 (2013).
37. Mills, M. C. & Rahal, C. A scientometric review of genome-wide association studies. *Commun. Bio.* **2**, 9 (2019).
38. Harden, K. P. *The Genetic Lottery: Why DNA Matters for Social Equality*. (Princeton University Press, 2021).
39. Rawls, J. *A theory of justice*. (Belknap Press of Harvard University Press, 1999).
40. Roemer, J. E. *Equality of Opportunity*. (Harvard University Press, 1998).
41. Gromet, D. M., Hartson, K. A. & Sherman, D. K. The politics of luck: Political ideology and the perceived relationship between luck and success. *J. Exp. Soc. Psychol.* **59**, 40–46 (2015).

42. Cappelen, A. W., Konow, J., Sørensen, E. Ø. & Tungodden, B. Just luck: An experimental study of risk-taking and fairness. *Am. Econ. Rev.* **103**, 1398–1413 (2013).
43. Almås, I., Cappelen, A. W., Sørensen, E. Ø. & Tungodden, B. Fairness and the development of inequality acceptance. *Science* **328**, 1176–1178 (2010).
44. Cappelen, A. W., Sørensen, E. Ø. & Tungodden, B. Responsibility for what? Fairness and individual responsibility. *Eur. Econ. Rev.* **54**, 429–441 (2010).
45. Alesina, A. & Ferrara, E. L. Preferences for redistribution in the land of opportunities. *Journal of Public Economics* **89**, 897–931 (2005).
46. Alesina, A., Stantcheva, S. & Teso, E. Intergenerational mobility and preferences for redistribution. *Am. Econ. Rev.* **108**, 521–554 (2018).
47. Nozick, R. *Anarchy, state, and utopia*. vol. 5038 (New York: Basic Books, 1974).
48. Joly, Y. *et al.* Establishing the International Genetic Discrimination Observatory. *Nat. Genet.* **52**, 466–468 (2020).
49. Benjamin, D. J. *et al.* The genetic architecture of economic and political preferences. *PNAS* **109**, 8026–8031 (2012).
50. van der Loos, M. J. H. M. *et al.* Candidate gene studies and the quest for the entrepreneurial gene. *Small Bus. Econ.* **37**, 269–275 (2011).
51. Beauchamp, J. P. *et al.* Molecular Genetics and Economics. *J. Econ. Perspect.* **25**, 57–82 (2011).
52. Karlsson Linnér, R. *et al.* Genome-wide association analyses of risk tolerance and risky behaviors in over 1 million individuals identify hundreds of loci and shared genetic influences. *Nat. Genet.* **51**, 245–257 (2019).
53. Hewitt, J. K. Editorial policy on candidate gene association and candidate gene-by-environment interaction studies of complex traits. *Behav. Genet.* **42**, 1–2 (2012).
54. Bioethics, N. C. on. *Genetics and Human Behavior: The Ethical Context*. (Nuffield Council on Bioethics, 2002).
55. Visscher, P. M., Hill, W. G. & Wray, N. R. Heritability in the genomics era--concepts and misconceptions. *Nat. Rev. Genet.* **9**, 255–266 (2008).
56. Taubman, P. The determinants of earnings: Genetics, family, and other environments: A study of white male twins. *Am. Econ. Rev.* **66**, 858–870 (1976).
57. Turkheimer, E. Three laws of behavior genetics and what they mean. *Curr. Dir. Psychol. Sci.* **9**, 160–164 (2000).
58. Wood, A. R. *et al.* Defining the role of common variation in the genomic and biological architecture of adult human height. *Nat. Genet.* **46**, 1173–1186 (2014).
59. Locke, A. E. *et al.* Genetic studies of body mass index yield new insights for obesity biology. *Nature* **518**, 197–206 (2015).
60. Yengo, L. *et al.* Meta-analysis of genome-wide association studies for height and body mass index in ~700,000 individuals of European ancestry. *Human Molecular Genetics* **27**, 3641–3649 (2018).
61. Jansen, I. E. *et al.* Genetic meta-analysis identifies 9 novel loci and functional pathways for Alzheimer's disease risk. *bioRxiv* 258533 (2018) doi:10.1101/258533.
62. Lambert, J. C. *et al.* Meta-analysis of 74,046 individuals identifies 11 new susceptibility loci for Alzheimer's disease. *Nat. Genet.* **45**, 1452–1458 (2013).
63. Ripke, S. *et al.* Biological insights from 108 schizophrenia-associated genetic loci. *Nature* **511**, 421–427 (2014).
64. Pardiñas, A. F. *et al.* Common schizophrenia alleles are enriched in mutation-intolerant genes and in regions under strong background selection. *Nat. Genet.* **50**, 381–389 (2018).
65. Barban, N. *et al.* Genome-wide analysis identifies 12 loci influencing human reproductive behavior. *Nat. Genet.* **48**, 1462–1472 (2016).
66. Okbay, A. *et al.* Genetic variants associated with subjective well-being, depressive symptoms, and

neuroticism identified through genome-wide analyses. *Nat. Genet.* **48**, 624–633 (2016).
67. Turley, P. *et al.* Multi-trait analysis of genome-wide association summary statistics using MTAG. *Nat. Genet.* **50**, 229–237 (2018).
68. Rietveld, C. A. *et al.* GWAS of 126,559 individuals identifies genetic variants associated with educational attainment. *Science* **340**, 1467–1471 (2013).
69. Lander, E. S. & Schork, N. J. Genetic dissection of complex traits. *Science* **265**, 2037–2048 (1994).
70. Thorgeirsson, T. E. *et al.* A variant associated with nicotine dependence, lung cancer and peripheral arterial disease. *Nature* **452**, 638–642 (2008).
71. Amos, C. I. *et al.* Genome-wide association scan of tag SNPs identifies a susceptibility locus for lung cancer at 15q25.1. *Nat. Genet.* **40**, 616–622 (2008).
72. Hung, R. J. *et al.* A susceptibility locus for lung cancer maps to nicotinic acetylcholine receptor subunit genes on 15q25. *Nature* **452**, 633–637 (2008).
73. Daetwyler, H. D., Villanueva, B. & Woolliams, J. A. Accuracy of predicting the genetic risk of disease using a genome-wide approach. *PLoS One* **3**, e3395 (2008).
74. de Vlaming, R. *et al.* Meta-GWAS accuracy and power (MetaGAP) calculator shows that hiding heritability is partially due to imperfect genetic correlations across studies. *PLoS Genet.* **13**, (2017).
75. Young, A. I., Benonisdottir, S., Przeworski, M. & Kong, A. Deconstructing the sources of genotype-phenotype associations in humans. *Science* **365**, 1396–1400 (2019).
76. Mostafavi, H. *et al.* Variable prediction accuracy of polygenic scores within an ancestry group. *Elife* **9**, (2020).
77. Winkler, T. W. *et al.* Quality control and conduct of genome-wide association meta-analyses. *Nat. Protoc.* **9**, 1192–1212 (2014).
